# Injury-induced CTE-like pathology emerges in a human multicellular *in vitro* brain model and reveals mitochondrial and neurovascular regulation

**DOI:** 10.64898/2025.12.29.696879

**Authors:** Sunghyun Jun, Daniel S Hinrichsen, Sahan BS Kansakar, Charitha C Anamala, Jada Teregeyo, Haiden C. Roberts, Colleen M Arrasmith, Mitchell D Thielen, Volha Liaudanskaya

**Affiliations:** Biomedical Engineering Department, University of Cincinnati, Cincinnati, USA; Neuroscience Graduate Program, University of Cincinnati, College of Medicine, Cincinnati, OH, USA

## Abstract

Chronic traumatic encephalopathy (CTE) is a progressive neurodegenerative disease linked to repetitive mild head impacts, but no human-based experimental system exists to study injury-induced CTE-like pathology. Here, we establish a long-lived, human multicellular *in vitro* brain platform in which controlled mechanical injury induces key cellular features of CTE-like pathology. Injured cultures developed persistent tau phosphorylation, axonal degeneration, chronic inflammation, and metabolic dysfunction without widespread neuronal loss, consistent with progressive pathology rather than acute toxicity. To assess physiological relevance, we integrated transcriptomic profiles from the model with postmortem human CTE brain datasets. This analysis revealed striking convergence at the level of disease-associated modules and pathways, with endothelial cells emerging as critical contributors to CTE-like transcriptional programs. Using this human-based system, we further identified delayed mitochondrial dysfunction as a prominent and sustained feature of injury-induced pathology. Together, these findings establish the first human *in vitro* platform for studying injury-induced CTE-like pathology and identify neurovascular and mitochondrial regulation as central components of chronic neurodegeneration following repetitive mild brain injury.

## Introduction

Chronic Traumatic Encephalopathy (CTE) is a progressive neurodegenerative disease linked to years of repetitive mild head impacts, most often affecting athletes, military personnel, and others exposed to cumulative brain trauma ^1–3^. Patients develop mood disturbances, cognitive decline, impulsivity, and eventually dementia, yet definitive diagnosis remains postmortem, and no disease-modifying therapies currently exist ^4,5^. Neuropathological studies consistently describe perivascular deposits of hyperphosphorylated tau, axonal and dendritic degeneration, chronic microgliosis, astrocytic reactivity, and, in subsets of cases, TDP-43 pathology ^6–8^. Despite these defining features, the biological mechanisms that connect survivable mild injuries to delayed and progressive pathology remain poorly understood ^8^.

Recent transcriptomic and neuropathological analyses of young athletes exposed to repetitive head impacts have begun to explain early disease-associated changes that precede widespread tau deposition ^9,10^. These studies reveal neuronal vulnerability, synaptic loss, microglial activation, astrocytosis, and pronounced vascular dysfunction, including endothelial stress and blood-brain barrier alterations, even in the absence of overt neurodegeneration ^7,11–15^. Together, these findings underscore that CTE is a fundamentally multicellular disorder involving coordinated dysfunction across neural and vascular compartments. However, because these observations derive from postmortem tissue, they provide only static snapshots of disease and cannot resolve how pathology evolves over time or which cellular components actively drive progression versus compensation.

To overcome this limitation, we established a long-lived, human-derived 3D brain culture platform that enables precise control of injury magnitude, frequency, and interval under survivable conditions. This system supports sustained co-culture of neurons, astrocytes, and microglia (tri-cultures) ^16^ and can be extended to incorporate endothelial cells and pericytes (penta-cultures) ^17^, reconstructing key elements of the human neurovascular unit ^16–24^. By applying controlled mechanical injury to genetically unmodified human cells, this platform enables longitudinal investigation of injury-induced multicellular responses that cannot be accessed in human tissue or short-lived culture systems. Importantly, it allows direct comparison of neural-only and neurovascular contexts under identical injury conditions.

Using this approach, we demonstrate that repetitive mild injury is sufficient to induce progressive, CTE-like pathology in a human *in vitro* system, including early tau phosphorylation, dendritic and synaptic degeneration, axonal injury, chronic neuroinflammation, perivascular stress, and delayed TDP-43 pathology ^7,11–15^. Importantly, injury-induced transcriptional programs in this system show striking convergence with postmortem human CTE brain datasets at the level of disease-associated modules and pathways, supporting the physiological relevance of the model ^9,25^.

Together, this human multicellular *in vitro* injury model establishes a previously unavailable experimental framework in which injury-induced CTE-like pathology emerges and identifies neurovascular interactions and mitochondrial dysfunction as central regulators of chronic neurodegeneration following repetitive mild brain injury.

## Results

### Mapping injury thresholds that permit survivable trauma for long-term modeling

We employed a previously established long-lived, human-derived 3D brain tissue platform that maintains stable neuronal, glial, and multicellular organization over extended culture periods and is amenable to controlled mechanical perturbation ^16,17,19,21–24^ (Fig.1A). This system enables systematic interrogation of injury parameters while preserving tissue viability, making it well-suited for defining thresholds that permit survivable trauma and delayed pathological evolution. To establish an *in vitro* model of CTE-like progression, we first sought to define the range of mechanical insults that cultures could withstand without immediate catastrophic damage, yet remain vulnerable to delayed pathology. We applied focal (1 mm tip, shear-dominant) or diffuse (3 mm tip, compressive) impacts at graded velocities (1 – 6m/s) and repeated up to three times, and assessed cellular responses 24 h after the final insult (Fig.1B-I).

**Fig. 1.**
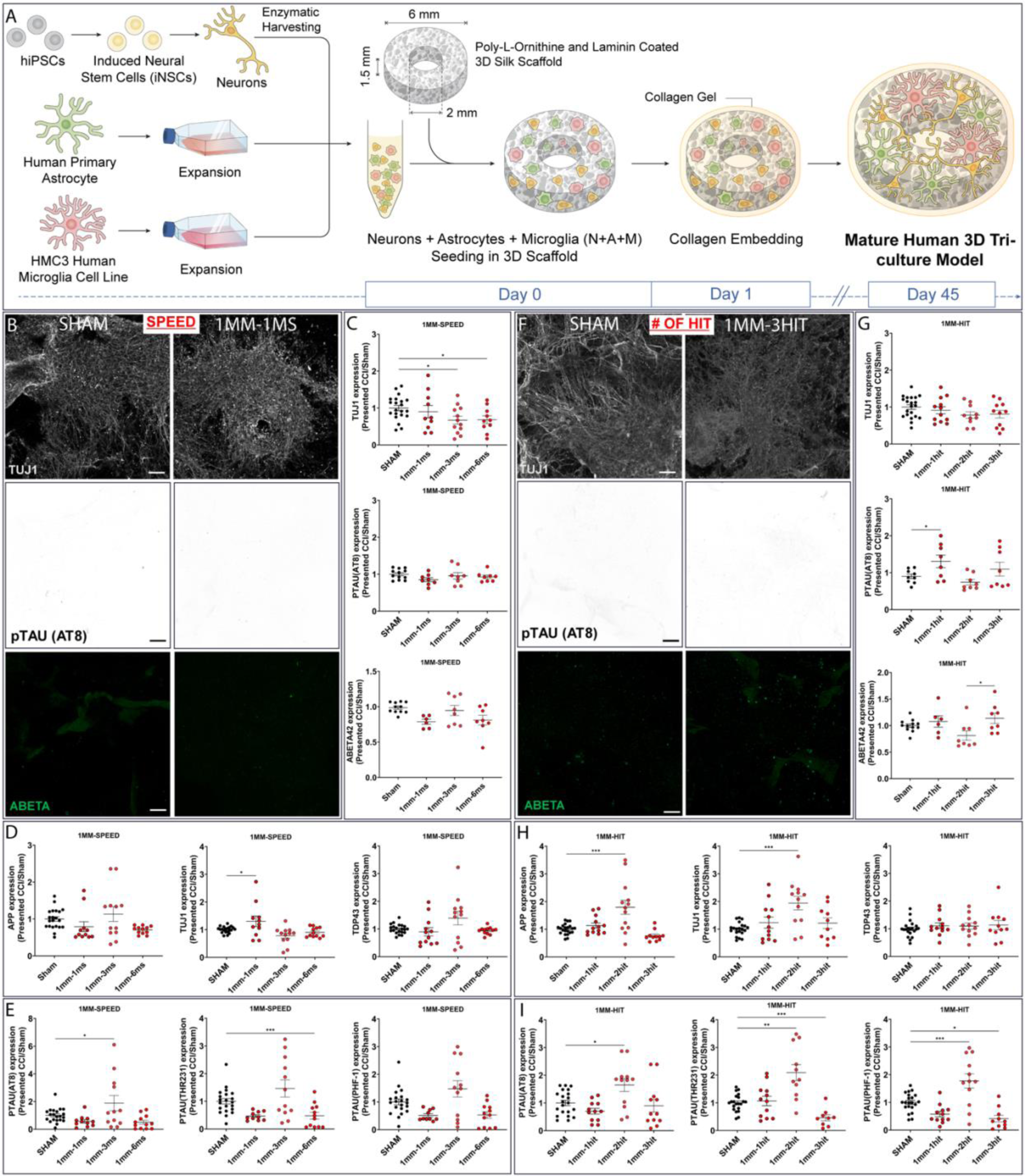
1 mm tip size injury area with 1 m/s speed of 3 hits is the mechanical force that is required to recapitulate clinically relevant injury threshold. Experiments were conducted to assess the effects of varying speeds (1 m/s, 3 m/s, and 6 m/s) and different injury counts (1 hit, 2 hits, and 3 hits) with a 1 mm tip size. **A** Schematic representation of the preparation of 3D tricultures. **B, F** Representative image of NAM tri-culture 1 mm tip size with various speeds, and 1 mm tip size and 1 m/s speed with various injury counts mentioned above. Immunofluorescent imaging of Tuj1 marker for neuronal networks; phosphorylated TAU(AT8) for degenerating neurons; and Ab42 for pathological proteinopathy. Scale bar: 100 µm. **C, G** Quantifications of the images from panel B, F (n = 6 – 10 scaffolds per condition). **D-H and E-I** Intracellular markers – Western Blot quantification of AD-related (APP, TDP-43), CTE-related (pTAU(AT8), pTAU(THR231), and pTAU(PHF-1)), and neuronal pan (Tuj1) markers (n = 9 – 12 scaffolds per condition) for both speed (D,E) and number of hits **(H,I**). Data presented mean ± SEM of n = 6 – 12 scaffolds per condition. *, **, *** indicates a significant difference with p<0.05, p<0.01, p<0.001, respectively. A one-way ANOVA (analysis of variance) test was used to determine the difference between the control and experimental groups. The ROUT outlier analysis method was used to exclude statistical outliers. Experiments were replicated at least three times, unless specified otherwise. Confocal images of pTau(AT8) and Ab42 were replicated twice.

Focal shear injuries (1 mm tip) revealed a narrow survivable window (Fig.1B-E). At low velocity (1 m/s), networks remained structurally intact, with no changes in tau phosphorylation or cytoskeletal markers. At moderate velocity (3 m/s), cultures began to show subtle stress responses: modest induction of early pTAU (AT8) and reduced TUJ1, without widespread collapse, suggesting the threshold at which pathology can be initiated while preserving network viability. At high velocity (6m/s), however, the system failed, with widespread neuronal loss and loss of pTAU signals, reflecting devastating damage that precludes further modeling. Repeated injuries further refined this window: two hits enhanced pathological signatures (AT8, Thr231, PHF-1, APP), whereas three hits paradoxically attenuated tau phosphorylation while preserving network structure, consistent with an acute shock response but overall survival (Fig.1F-I). These findings defined the 1 mm / 3-hit paradigm as the lowest intensity regime that maintains tissue viability while seeding delayed pathology.

Diffuse compressive injuries (3 mm tip) produced a different profile (fig.S1A-C, S2D-F). Shown in fig.S1A-C, even a single low-velocity hit (1 m/s) was sufficient to activate widespread pathology, with significant induction of AT8, Thr231, and PHF-1, suggesting that compressive forces more efficiently trigger injury responses. At intermediate velocity (3 m/s), tau signals were absent, but TDP-43 was strongly induced, pointing to a force-dependent shift between degenerative programs. High-intensity injuries (6 m/s) caused catastrophic network loss, preventing long-term study. With repetition shown in fig.2D-F, the dose dependence became clear: one hit remained largely sub-threshold, two hits produced APP suppression without tau induction, and three hits combined APP loss with late-stage tauopathy (PHF-1) and TUJ1 decline.

**Fig. 2.**
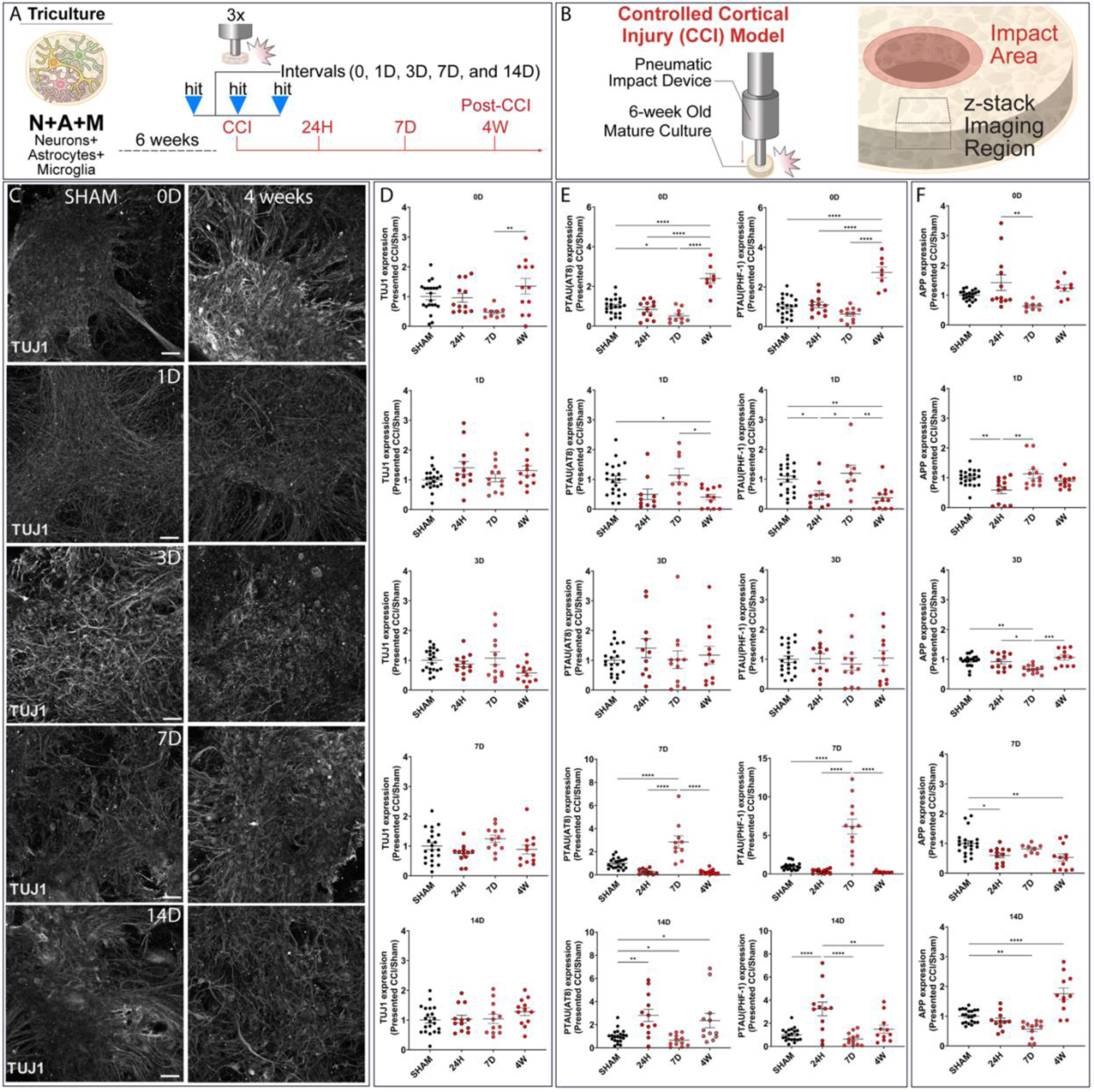
A 14-day interval serves as the primary driving force behind the onset and progression of Taupathy. Three mild repetitive injuries (1 mm-1 m/s) were applied to the NAM triculture model with various intervals (0-day, 1-day, 3-day, 7-day, and 14-day intervals). **A** Schematic representation of injury experimental design. **B** Contusion (Controlled cortical impact model (CCI)) injury and schematic representation of the injury area, and image acquisition. **C** Representative Tuj1 (pan neuronal marker) staining of sham and 4 weeks post-injury condition (24 hours and 7 days post-injury condition in fig.S3). Scale bar: 100 µm. **D** Quantifications of Tuj1 neuronal networks. **E, F** Intracellular markers – Western Blot quantification of AD-related (APP) and CTE-related (pTAU(AT8) and pTAU(PHF-1)) markers. Data presented mean ± SEM of n = 9 – 12 scaffolds per condition. **\*,** **, **** indicates a significant difference with p<0.05, p<0.01, p<0.0001, respectively. A one-way ANOVA (analysis of variance) test was used to determine the difference between the control and experimental groups. The ROUT outlier analysis method was used to exclude statistical outliers. Experiments were replicated at least three times.

Together, these results define the injury parameters that cultures can survive while remaining vulnerable to disease. Focal shear required moderate repetition to generate delayed pathology without overt neuronal loss, whereas diffuse compression was a more efficient trigger but less suited for long-term modeling because even low-level insults caused immediate tau pathology. This mapping establishes the clinically relevant injury window necessary for modeling how repetitive mild trauma evolves into CTE-like disease.

### The injury interval determines whether repetitive trauma allows for recovery or results in neurodegenerative-like pathology

Having defined the acute thresholds for survivable injury, we next examined how the timing between repetitive insults influences the long-term trajectory of pathology (Fig.2A-B). Three mild impacts were delivered at intervals of 0, 1, 3, 7, or 14 days, and cultures were analyzed at 24 h, 7 days, and 4 weeks (Fig.2A-F, fig.S3-5).

Short intervals (0–3 days) produced minimal or reversible effects, with cultures largely maintaining neuronal Tuj1 viability (Fig.2C-D, fig.S3). Consecutive hits (0-day interval) caused only transient structural disruption, marked by early MAP2 and SYN1 (shown in fig.S4) elevation at 24 h and a temporary decrease in AT8 at 7 days, followed by recovery of tau signals (AT8 and PHF-1) at 4 weeks (Fig.2E). One-day and three-day intervals similarly produced little long-term pathology, aside from a temporary MAP2 dip at 7 days in the 3-day condition, which normalized by 4 weeks (fig.S4). These findings indicate that closely spaced injuries tend to overwhelm the system transiently but do not establish sustained degeneration.

A 7-day interval promoted structural resilience. Cultures mounted a coordinated stress–recovery response, with transient tau phosphorylation (AT8 and PHF-1) at 7 days that resolved completely by 4 weeks, accompanied by significant increases in TUJ1 and MAP2, suggesting cytoskeletal reinforcement (Fig.2C-F, fig.S3-4). This interval provided sufficient time for intrinsic repair mechanisms to restore stability, preventing progression to chronic pathology.

By contrast, a 14-day interval sensitized the system to delayed degeneration. Cultures displayed an early, exaggerated phosphorylation response (AT8 and PHF-1) at 24 h, which subsided by 7 days (Fig.2C-F, fig.S3-4). However, by 4 weeks, they transitioned into a chronic disease-like state, characterized by persistent AT8 elevation and APP accumulation, despite normalization of PHF-1. Thus, the extended recovery period did not confer protection but instead primed the tissue for maladaptive responses and long-term vulnerability.

Together, these findings show that the interval between injuries is a critical determinant of whether cultures resolve damage or progress toward chronic pathology. While short intervals favor transient disruption and a 7-day interval supports recovery, longer spacing (14 days) leaves the system hypersensitive, converting otherwise survivable injuries into a delayed degenerative cascade.

### Repetitive mild injury induces a staged transition from synaptic crisis to chronic taupathy

After establishing survivable injury parameters, we next examined the temporal evolution of pathology over an extended 8-week period following repetitive mild injury (Fig. 3A–E, Fig. S6). To capture both intracellular and extracellular dynamics, we combined immunofluorescence imaging, Western blot analysis, and multiplex neuropanel measurements. This integrated approach revealed a structured progression in which cultures initially survived injury but gradually transitioned into a chronic, molecularly defined degenerative state.

**Fig. 3.**
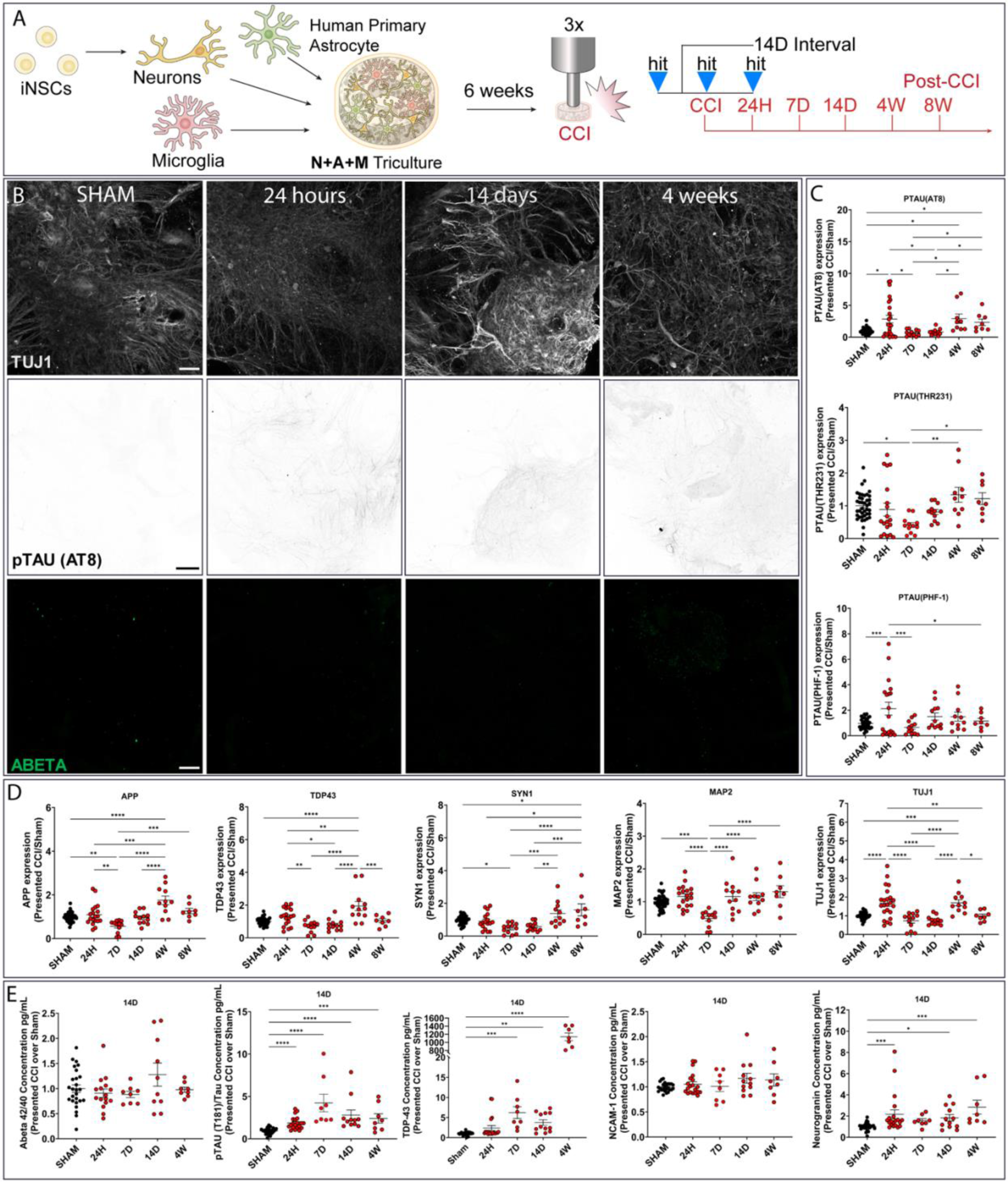
Early synaptic disruption precedes delayed taupathy after repetitive mild injury. Three mild repetitive injuries (1 mm-1 m/s) were applied to the NAM-triculture model with a 14-day interval, incorporating extended time points at 24 hours, 7 days, 14 days, 4 weeks, and 8 weeks. **A** Schematic representation of injury experimental design. **B** Representative Tuj1 (pan neuronal marker) immunofluorescent staining of sham, 24 hours, 14 days, and 4 weeks post-injury condition; phosphorylated TAU(AT8) for degenerating neurons; and Ab42 for pathological proteinopathy. Quantifications of representative images are shown in fig.S6. Scale bar: 100 µm. **C, D** Intracellular markers – Western Blot quantification of AD-related (APP, TDP-43), CTE-related (pTAU(AT8), pTAU(THR231), and pTAU(PHF-1)), and Neuronal structure and Synaptic health (Tuj1, MAP2, and SYN1) markers. **E** Extracellular markers were assessed with a 9-plex neurodegeneration panel at different post-injury time points (24H, 72H, 1W, and 4W). Extracellular markers: Amyloid beta 1-42 presented over Amyloid beta 1-40, phosphorylated TAU (pT181) over TAU, TDP-43, NCAM-1, and Neurogranin. Data presented mean ± SEM of n = 6 – 12 scaffolds per condition. **\*,** **, ***, **** indicates a significant difference with p<0.05, p<0.01, p<0.001 p<0.0001, respectively. A one-way ANOVA (analysis of variance) test was used to determine the difference between the control and experimental groups. Brown-Forsythe and Welch ANOVA test was used for TDP-43 and Abeta42 from extracellular markers. The ROUT outlier analysis method was used to exclude statistical outliers. Experiments were replicated at least three times, unless specified otherwise. Intracellular markers for the 8W time point and extracellular markers for 7D and 4W time points were replicated two times.

During the first week after the final injury, pathology was dominated by synaptic and dendritic dysfunction rather than overt cell loss. TUJ1 networks appeared fragmented by immunofluorescence, and structural proteins MAP2 and SYN1 declined significantly by 7 days, consistent with early dendritic and synaptic destabilization (Fig. 3B–D). Neurogranin release increased acutely at 24 h, further supporting early dendritic stress (Fig. 3E). In parallel, tau pathology was rapidly engaged: intracellular AT8 phosphorylation increased within 24 h (Fig. 3C), and extracellular pTAU/TAU ratios peaked by 7 days (Fig. 3E). Together, these findings indicate that the earliest response to survivable repetitive injury is a synaptic–dendritic crisis accompanied by early tau phosphorylation, rather than catastrophic tissue collapse.

Between 14 days and 4 weeks, cultures progressed into a degenerative phase despite partial preservation of network structure. MAP2 levels showed partial recovery, yet intracellular APP and TDP-43 accumulated significantly by 4 weeks (Fig. 3D). Extracellular TDP-43 levels increased in parallel, indicating both intracellular accumulation and pathological release. Notably, the Aβ42/40 ratio remained unchanged throughout the time course, distinguishing this trajectory from Alzheimer-like pathology and supporting alignment with CTE-specific processes (Fig. 3E, Fig. S6). Thus, cultures remained structurally viable while developing a sustained molecular signature of neurodegeneration.

By 8 weeks, several acute injury markers had resolved, with APP and TDP-43 returning toward baseline. In contrast, AT8 phosphorylation remained persistently elevated, defining a stable plateau of early-stage tauopathy (Fig. 3C–D). SYN1 levels increased significantly at this late stage, suggesting maladaptive synaptic remodeling rather than restoration of normal connectivity. Although overall network viability was maintained, neuronal architecture remained disorganized, consistent with a long-lived but dysfunctional state.

Collectively, these results demonstrate that repetitive mild injury initiates a delayed and staged degenerative process rather than immediate tissue failure. An initial phase of synaptic and dendritic disruption gives way to tau- and TDP-43–associated pathology, followed by stabilization into a chronic AT8-positive tauopathic state with persistent structural disorganization. This trajectory mirrors clinical observations in CTE, where apparently recoverable injuries precede the gradual emergence of long-term neurodegeneration ^11,14,26^.

### The neurovascular unit reshapes the temporal cascade of pathology in penta-cultures

Because perivascular tau deposition is a defining feature of human CTE, we next asked how the neurovascular unit (NVU) alters the trajectory of disease. To test this, we incorporated endothelial cells and pericytes into our cultures to generate penta-cultures and subjected them to the 14-day interval injury paradigm ^17^ (Fig.4A-F, fig.S7). The presence of the NVU did not prevent pathology entirely but fundamentally reshaped its timing and character, shifting the system from immediate collapse toward delayed, transient degenerative episodes.

**Fig. 4.**
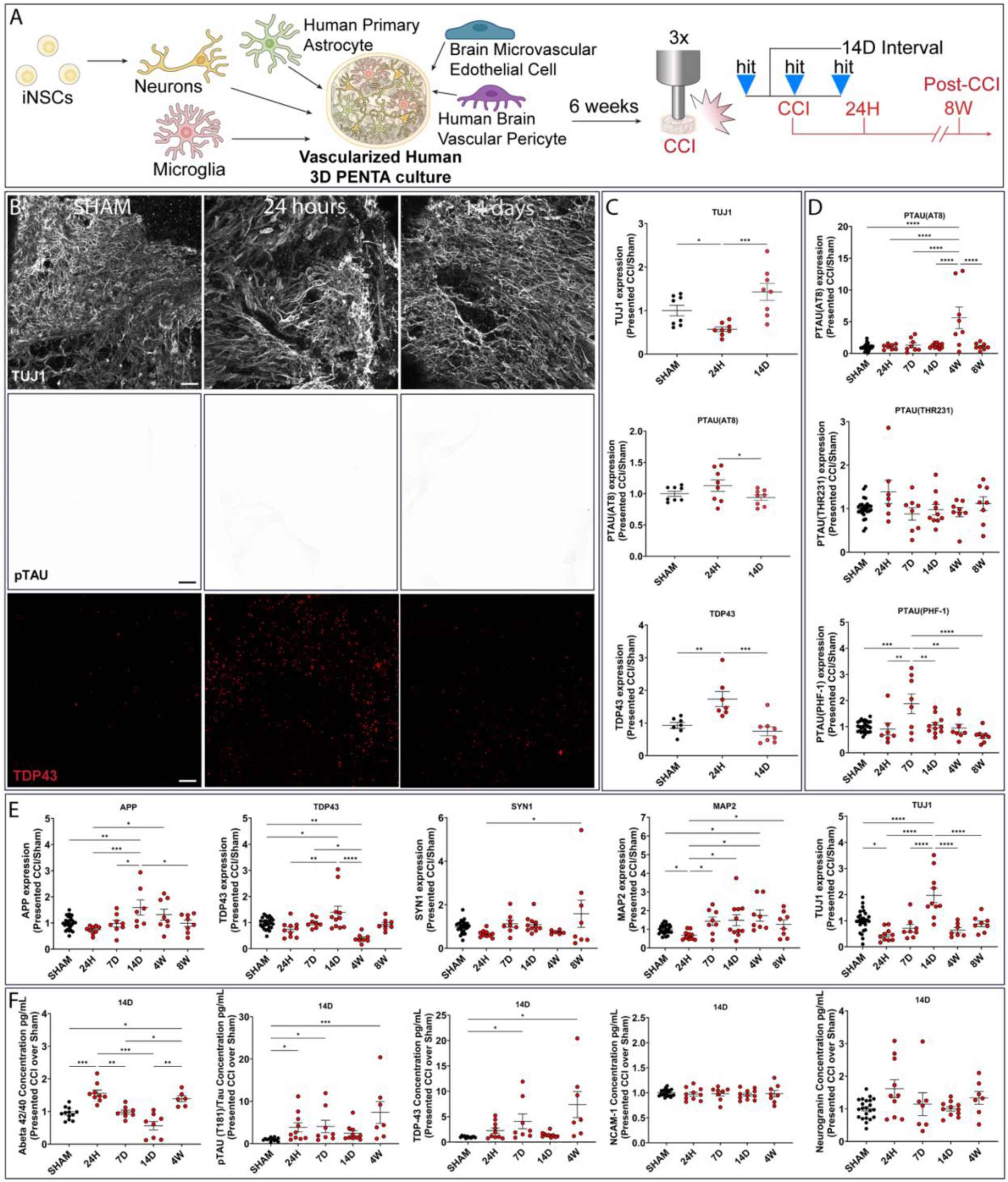
The neurovascular unit reshapes the temporal trajectory of injury-induced neurodegeneration. Three mild repetitive injuries (1mm-1m/s) were applied to the vascularized PENTA-triculture model with a 14-day interval, incorporating extended time points at 24 hours, 7 days, 14 days, 4 weeks, and 8 weeks. **A** Schematic representation of injury experimental design for penta-culture model. **B** Representative image of PENTA sham, PENTA 24 hours post-injury, and PENTA 14 days post-injury. Immunofluorescent imaging of Tuj1 marker for neuronal networks; phosphorylated TAU(AT8) for degenerating neurons; and TDP-43 for pathological proteinopathy. Scale bar: 100 µm. **C** Quantifications of Tuj1 neuronal networks, pTAU(AT8) aggregates, and TDP-43 at different post-injury time points. **D, E** Intracellular markers – Western Blot quantification of AD-related (APP, TDP-43), CTE-related (pTAU(AT8), pTAU(THR231), and pTAU(PHF-1)), and Neuronal structure-Synaptic health (Tuj1, MAP2, and SYN1) markers. **F** Extracellular markers were assessed with a 9-plex neurodegeneration panel at different post-injury time points (24H, 72H, 1W, and 4W). Extracellular markers: Amyloid beta 1-42 presented over Amyloid beta 1-40, phosphorylated TAU (pT181) over TAU, TDP-43, NCAM-1, and Neurogranin. Data presented mean ± SEM of n = 6 – 12 scaffolds per condition. **\*,** **, ***, **** indicates a significant difference with p<0.05, p<0.01, p<0.001 p<0.0001, respectively. A one-way ANOVA (analysis of variance) test was used to determine the difference between the control and experimental groups. Brown-Forsythe and Welch ANOVA test was used for Neurogranin and Abeta42, and Kruskal-Wallis test was used for TDP-43 from extracellular markers. The ROUT outlier analysis method was used to exclude statistical outliers. Experiments were replicated at least three times, unless specified otherwise. Confocal images of Tuj1, pTAU(AT8), and TDP-43 were replicated two times. Intracellular markers for the 1W, 4W, and 8W time points and extracellular markers for 7D and 4W time points were replicated two times.

Within the first 24 hours, neuronal networks displayed significant structural stress, with reduced TUJ1 and MAP2 expression and visible fragmentation of neurites (Fig.4B-E). However, unlike NAM cultures, tau phosphorylation remained at baseline (AT8 negative), indicating that the NVU buffered the system from immediate tauopathy despite clear mechanical damage (Fig.4C-D). By 7 days, a transient rise in late-stage tau (PHF-1) was detected, but early-stage AT8 remained absent. These findings suggest that vascular elements absorb or redistribute early injury signals, preventing premature activation of core tau pathology.

By 14 days, structural integrity was largely restored. TUJ1 expression rebounded, and neurite density appeared comparable to sham conditions. To directly assess vascular integrity during these early phases, we quantified endothelial network architecture in PENTA cultures following injury (fig. S7). Confocal imaging of hBMEC-derived GFP revealed pronounced but non-destructive remodeling of vascular-like networks after repetitive mild injury. Quantitative AngioTool analysis demonstrated a significant reduction in junction density at 24 hours, accompanied by increased vessel lacunarity and vessel length at both 24 hours and 14 days compared to sham controls. These changes are consistent with early vascular destabilization and increased network heterogeneity rather than overt vessel loss. Importantly, despite these architectural alterations, endothelial networks remained continuous and viable across time points, indicating that vascular elements persist through injury and recovery phases. This pattern supports the concept that the neurovascular unit undergoes structural reorganization in response to mechanical trauma, transiently compromising network compactness while preserving overall vessel continuity. Such remodeling likely contributes to the buffering of acute neuronal injury observed in PENTA cultures, while simultaneously creating a state of vascular stress that precedes the delayed emergence of tau and TDP-43 pathology.

Extracellular stress markers also normalized, with both secreted pTAU and TDP-43 returning to baseline. Despite this apparent recovery, intracellular pathology accumulated silently: APP and TDP-43 were both significantly elevated, consistent with toxic substrate buildup within neurons and glia (Fig.4E). Immunofluorescence confirmed widespread TDP-43 puncta, revealing that the NVU sustained structural integrity while concealing emerging molecular pathology.

By 4 weeks, this compensatory state failed. AT8 tauopathy emerged robustly within cells, accompanied by significant extracellular release of pTAU and TDP-43 (Fig. 4D-E). At the same time, vascular stress resurfaced, evidenced by renewed increases in the Aβ42/40 ratio, linking endothelial dysfunction to the onset of tau and TDP-43 pathology (Fig. 4F). This phase marked the critical inflection point when NVU buffering was no longer sufficient, and the system transitioned into a full CTE-like degenerative state.

By 8 weeks, AT8 tau returned to sham levels, indicating that tauopathy in penta-cultures was transient rather than chronic. Extracellular markers of synaptic and dendritic integrity (NCAM1, Neurogranin, Synapsin) remained stable throughout, demonstrating that the NVU preserved long-term network health despite transient episodes of molecular pathology (Fig. 4F).

Together, these results demonstrate that the NVU delays, reshapes, and partially resolves the trajectory of injury-induced pathology. Rather than immediate tau-driven degeneration, penta-cultures pass through phases of buffered structural collapse, apparent recovery with intracellular accumulation, a delayed tau/TDP-43 crisis, and eventual resolution. This dual role, protective in the short term but stress-amplifying at critical points, mirrors the perivascular signatures of human CTE and highlights vascular elements as key modulators of disease fate ^11,27–29^.

### Injury-induced transcriptional programs in human brain injury are recapitulated at the pathway level in a multicellular *in vitro* model

To assess whether injury-induced transcriptional responses in our *in vitro* system recapitulate molecular features of human chronic traumatic encephalopathy (CTE), we integrated published postmortem human brain transcriptomic datasets, including bulk and cell-type-resolved RNA sequencing from individuals with CTE pathology ^9,10,25^. Human datasets were used to define disease-associated gene co-expression modules and pathway signatures, which were subsequently projected onto transcriptomic profiles from NAM and PENTA cultures following injury. This framework enabled direct comparison of global transcriptional organization across systems using unbiased, module-based metrics rather than selected gene lists.

Using all expressed genes assigned to each human-defined module, we evaluated module-trait relationships to determine whether coordinated transcriptional programs associated with human CTE were preserved *in vitro* (Fig. 5A-B, fig. S8). Twelve human CTE-associated modules were analyzed at early (24 h) and late (8 w) post-injury time points. At 24 h, nine of twelve modules in PENTA cultures showed concordant directionality with human CTE traits, whereas NAM cultures exhibited concordance in five modules. At 8 w, eight of twelve modules in PENTA cultures remained directionally aligned with human CTE, while NAM cultures showed concordant regulation in six modules. Across both time points, PENTA cultures displayed stronger and more consistent module-trait correlations than NAM cultures.

**Fig. 5.**
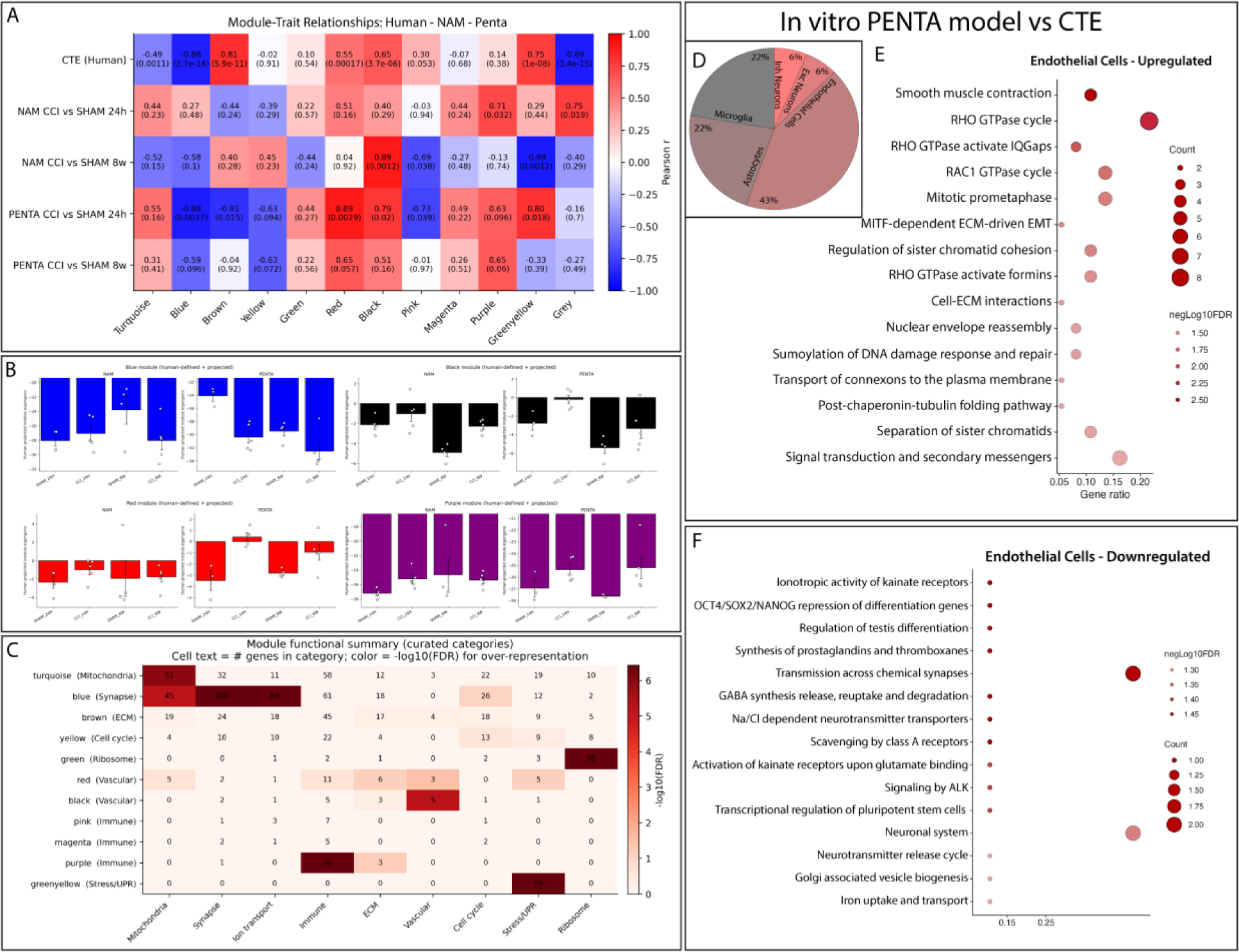
Human relevance of *in vitro* injury responses and pathway-level convergence with CTE. Human transcriptomic datasets from chronic traumatic encephalopathy (CTE) were integrated with multicellular human brain injury models to assess cross-system transcriptional alignment at the gene, module, and pathway levels. **A** Proportional contribution of major neural and vascular cell types to human CTE-associated transcriptional signatures derived from published postmortem CTE datasets ^9,25^ and mapped to corresponding cell types. **B-C** Human-defined co-expression modules were projected onto NAM and PENTA transcriptomes using all expressed genes assigned to each module. Module eigengene expression is shown for sham and injured conditions at matched time points, enabling comparison of coordinated transcriptional program engagement across systems. For each module, pathway activity scores are shown for time-matched sham and injured samples, with bars extending upward indicating relative enrichment of upregulated pathways and bars extending downward indicating enrichment of downregulated pathways compared to sham controls. Because pathway scores reflect absolute expression-derived values, sham samples are not constrained to a fixed reference level and capture baseline temporal dynamics of the system. **D-E** Pathway-level enrichment analyses were performed using injury-associated transcriptional changes *in vitro* without restricting input to concordant gene subsets.

In PENTA cultures, multiple modules exhibited coordinated regulation that matched the direction and relative magnitude of changes observed in human CTE datasets across time (Fig. 5B-C; fig. S8), including modules related to mitochondrial and metabolic regulation, ion transport, cytoskeletal organization, and inflammatory signaling. In contrast, NAM cultures showed smaller effect sizes and less consistent alignment across time for these modules, although directional concordance increased at 8 weeks relative to early time points.

Several modules displayed discordant behavior between *in vitro* systems and human datasets (fig. S8). These modules were enriched for pathways related to acute stress responses or transient injury-associated signaling that were prominent at early *in vitro* time points but not sustained in chronic human pathology (Fig.5C). Such discordant modules were more frequently observed in NAM cultures than in PENTA cultures.

Next, to identify cellular contributors to these module-level similarities, we compared bulk PENTA transcriptomes with cell-type-resolved RNA sequencing data from human RHI and RHI+CTE brains (Fig. 5D-F; fig. S9). Endothelial cells showed the strongest alignment between PENTA cultures and human disease datasets and represented the largest divergence between PENTA and NAM systems. In both PENTA cultures and human CTE datasets, endothelial cells exhibited the suppression of metabolic and barrier-associated programs, accompanied by the induction of cytoskeletal remodeling, inflammatory signaling, and stress-response pathways (Fig. 5E-F). These endothelial-associated transcriptional changes were substantially attenuated or absent in NAM cultures.

Additional cell types showed conserved but distinct transcriptional responses. Astrocytes demonstrated reduced expression of synaptic support and homeostatic pathways alongside induction of inflammatory and stress-associated programs (fig. S9). Microglia exhibited upregulation of immune activation, interferon signaling, and cytoskeletal remodeling pathways following injury. Neuronal populations exhibited suppression of synaptic organization and ion transport pathways, accompanied by concurrent activation of stress- and cell-cycle-associated programs. While these responses were detected in both NAM and PENTA cultures, their directional alignment and pathway composition were more consistent with human disease signatures in PENTA cultures.

Together, these analyses demonstrate that PENTA cultures preserve a greater fraction of human CTE-associated transcriptional modules and pathway-level organization than NAM cultures across early and late post-injury time points, with particularly strong alignment at the endothelial and mitochondrial-associated module levels.

### Cell-specific mitochondrial programs dictate resilience versus chronic dysfunction

Building on the pathway- and module-level analyses in Fig. 5, which identified mitochondrial-associated programs as a dominant axis of concordance between PENTA cultures and human CTE, we next examined whether these transcriptional signatures translated into functional differences in cellular bioenergetics. We therefore performed longitudinal metabolic profiling to directly test how mitochondrial activity evolves after repetitive mild trauma in NAM and NVU-containing PENTA cultures (Fig. 6A-C).

**Fig. 6.**
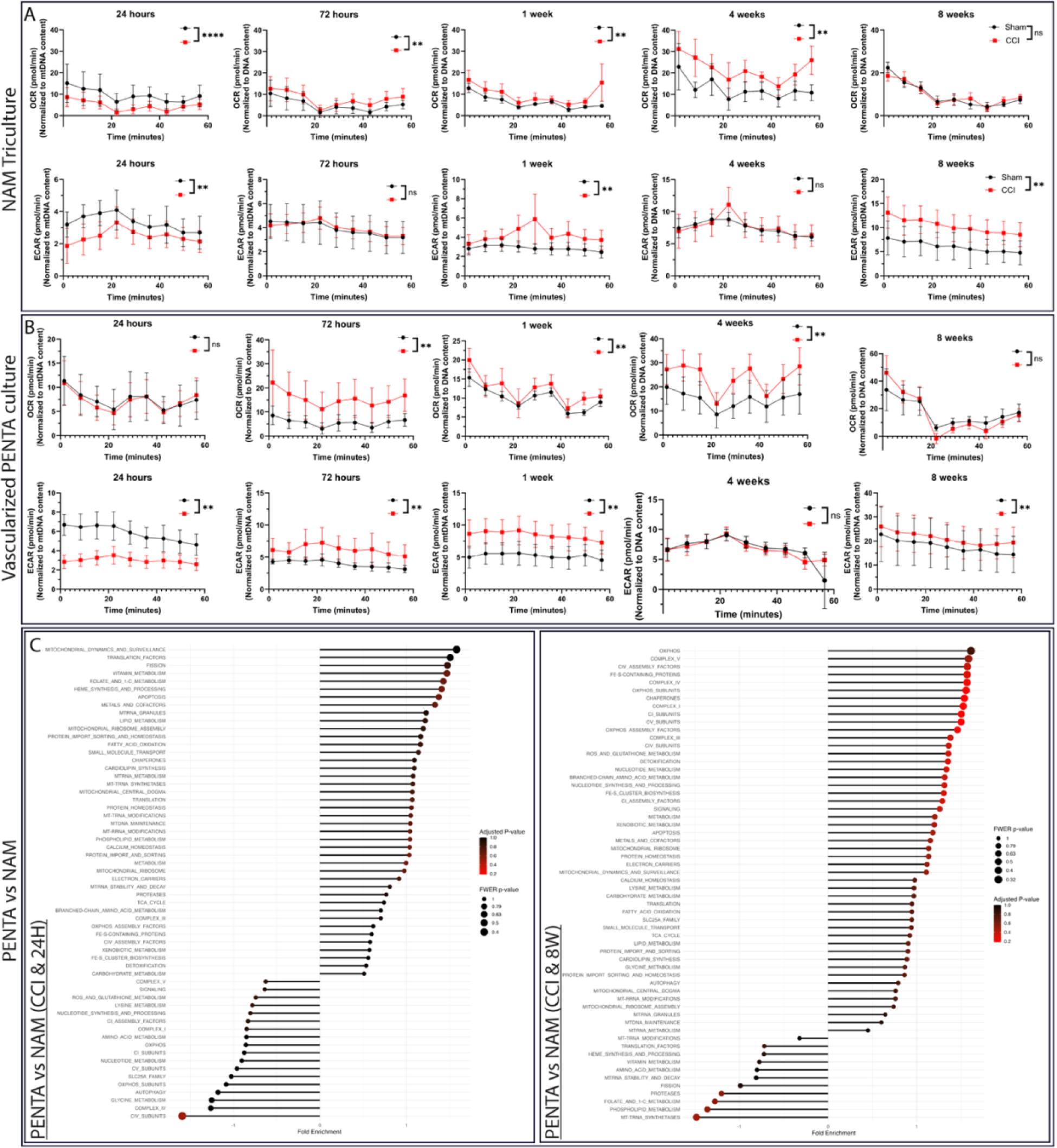
Divergent mitochondrial bioenergetic trajectories distinguish resilience from chronic dysfunction after injury. **A, B** NAM Triculture (**A**) and vascularized PENTA culture (**B**) ATP Rate Assay results comparing OCR and ECAR between different conditions across selected time points. Data presented are mean ± SEM of two independent experiments with a total of n=5 scaffolds for the sham groups and n=8 scaffolds for the injury groups across selected time points. **, **** indicates a significant difference with p<0.01, p<0.0001, respectively. Wilcoxon test was used to determine the difference between the control and experimental groups. The ROUT outlier analysis method was used to exclude statistical outliers. Presented OCR and ECAR rates were normalized to mitochondrial DNA content. **C** Differentially expressed genes (DEGs). Injury groups of PENTA 24H vs NAM 24H and PENTA 8W vs NAM 8W. Data presented are mean ± SEM of the experiment with n=4 scaffolds for the sham groups and n=5 scaffolds for the injury groups across selected time points for both PENTA and NAM.

In neuron-astrocyte-microglia tri-cultures, injury triggered an immediate and catastrophic collapse of both oxidative phosphorylation OCR and glycolysis ECAR within 24 hours (Fig. 6A). By one week, cultures rebounded into a hypermetabolic state, with OCR and ECAR both significantly elevated alongside increased TOMM20 expression (fig.S10), consistent with compensatory mitochondrial biogenesis. However, this response failed to produce durable adaptation. At four weeks, OCR remained elevated while TOMM20 normalized, indicating persistent mitochondrial overactivation without structural reinforcement. By eight weeks, OCR returned to baseline, but ECAR remained elevated, defining a stable state of glycolytic dependence. Transcriptomic analyses paralleled these functional dynamics: induction of mitochondrial maintenance programs, including ribosome assembly, chaperone activity, and mtDNA stability, was coupled to suppression of the TCA cycle, oxidative phosphorylation subunits, fatty acid oxidation, and mitophagy pathways (Fig. 6C, fig. S11). This pattern reflects a state of metabolic arrest that ultimately transitioned into inflammation-linked decline, locking tri-cultures into a chronically inefficient glycolytic phenotype ^16,30–32^.

In contrast, NVU-containing PENTA cultures exhibited a fundamentally different mitochondrial trajectory (Fig.6B-C, fig. S12). At 24 hours, OCR was preserved, and ECAR suppressed, indicating that vascular elements buffered the immediate metabolic collapse observed in tri-cultures (Fig. 6B). By one week, OCR and ECAR rose in parallel, reflecting a controlled hypermetabolic response rather than compensatory overdrive. At four weeks, OCR increased further and was accompanied by significant upregulation of TOMM20 (fig. S10), consistent with coordinated mitochondrial biogenesis and structural adaptation. By eight weeks, OCR normalized while ECAR remained modestly elevated, resembling the metabolic endpoint of tri-cultures but following a delayed, biphasic trajectory that avoided early collapse. Transcriptomic profiling supported this functional resilience: PENTA cultures activated mitochondrial fission, transport, and chaperone programs early, followed by re-establishment of oxidative phosphorylation subunits, assembly factors, and cofactor metabolism by eight weeks. Mitochondrial Complex II was preserved in PENTA cultures but suppressed in tri-cultures, highlighting NVU-mediated protection against apoptosis-associated mitochondrial failure.

Together, these findings provide functional validation of the mitochondrial pathways identified in Fig. 5 and establish mitochondria as decisive, cell-context-dependent regulators of injury outcome. In the absence of vascular support, neurons and glia undergo early mitochondrial collapse followed by maladaptive metabolic compensation that culminates in chronic glycolytic dysfunction. In contrast, inclusion of the neurovascular unit reshapes this trajectory by preventing acute mitochondrial failure, enforcing a phase of controlled biogenesis, and ultimately restoring oxidative capacity.

## Discussion

CTE pathology has been defined primarily through postmortem observations, including perivascular hyperphosphorylated tau, axonal injury, microgliosis, astrocytosis, and vascular abnormalities ^7,11–13,33^. Recent molecular and neuropathological profiling of young individuals exposed to repetitive head impacts further indicates that glial activation and endothelial stress can emerge early, before widespread neurodegeneration, supporting a multicellular disease process rather than a purely neuronal disorder ^9,34,35^. However, postmortem human tissue provides only static snapshots and cannot resolve how survivable injuries evolve into delayed and progressive pathology or which cellular compartments actively drive progression versus compensation ^5,7,36^. Here, we establish a one-of-a-kind human multicellular *in vitro* injury model that reproduces key cellular hallmarks of CTE-like pathology after controlled, survivable repetitive mild impacts, enabling mechanistic dissection with time-resolved sampling in a human-derived system ^16,19,20,24^.

Several features of our *in vitro* pathology align closely with human CTE. Cultures developed early AT8-positive tau phosphorylation, axonal injury (APP accumulation), and progressive dendritic/synaptic disruption without immediate catastrophic neuronal loss, consistent with the concept that early disease involves network destabilization and axonal pathology rather than frank tissue destruction ^7,11,12,33,37^. We also observed TDP-43-associated pathology emerging during the degenerative trajectory, consistent with reports that subsets of CTE cases exhibit broader proteinopathy beyond tau ^11,38^. Importantly, these phenotypes arose without genetic manipulation, indicating that repetitive survivable injury alone can be sufficient to trigger a delayed, multicellular degenerative cascade in human-derived tissue.

A central advance of this work is the demonstration of striking convergence between the *in vitro* injury response and human brain datasets at the module and pathway level, supporting physiological relevance beyond marker-based phenotyping ^34,35,39^. By projecting human CTE-associated transcriptional programs onto NAM and PENTA cultures, we found that vascularized PENTA cultures preserved a larger fraction of human CTE-linked transcriptional organization than NAM tri-cultures across acute and chronic timepoints, with particularly strong alignment in endothelial-associated programs ^7,34,35,39^. This result provides an experimental complement to human observations, highlighting vascular dysfunction and endothelial stress signatures in RHI/CTE-relevant contexts, and it supports the neurovascular unit as an active regulator of CTE-like progression rather than a secondary bystander ^7,9,11,33^.

The model also reveals mechanisms that cannot be directly inferred from postmortem human data: cell-context-dependent mitochondrial trajectories that segregate vulnerability from partial recovery. In NAM tri-cultures, repetitive injury produced an early collapse of oxidative phosphorylation and glycolysis, followed by a maladaptive rebound and eventual stabilization in a glycolysis-dependent state. In contrast, NVU-containing PENTA cultures buffered early metabolic failure, engaged delayed mitochondrial biogenesis, and more effectively restored oxidative programs over time. These findings extend prior evidence that mitochondrial dysfunction is a key feature of brain injury biology and can shape long-term outcomes ^4,16,36,40–43^. Together with the human convergence analyses, they support mitochondria not only as markers of injury but as active regulators of whether injury resolves, collapses, or progresses toward sustained pathology.

We further propose that delayed mitochondrial dysfunction may contribute to chronic pathology through metabolic-epigenetic coupling. Mitochondria supply metabolites that influence chromatin regulation and cellular repair capacity, including NAD⁺, acetyl-CoA, and α-ketoglutarate; persistent impairment in these pathways could create a permissive state for maladaptive transcriptional reprogramming after survivable injury ^22,39,44–46^. This framework is consistent with the broader concept that prolonged stress states, rather than injury exposure alone, may be required to shift tissues into progressive neurodegeneration, and it provides a mechanistic direction that can now be tested experimentally in a human system ^5,10,36,39^.

Beyond injury magnitude, our data indicate that injury timing is a major determinant of long-term outcome. We observed a non-linear response in which an intermediate injury pattern produced a more pronounced and sustained pathological trajectory than either fewer or more closely spaced impacts. This suggests that CTE-like progression is not governed solely by cumulative burden but can depend on whether a subsequent injury arrives during a vulnerable recovery window, when synaptic stability, mitochondrial reserve capacity, and neurovascular homeostasis are compromised but not irreversibly lost ^5,36,47,48^. Mechanistically, a first injury may initiate subthreshold stress that does not fully resolve; a second insult during this sensitization period may then lock the tissue into a chronic, partially compensated state that sustains inflammatory signaling and promotes delayed tau-associated degeneration. In contrast, very closely spaced or excessive injuries may trigger acute shutdown responses or catastrophic damage that shorten the window for slow disease-like evolution, yielding a qualitatively different trajectory.

The ability to compare NAM and PENTA systems under matched injuries also provides a direct experimental basis for a key human observation: vascular involvement is not merely correlative. In PENTA cultures, endothelial networks persisted through injury and underwent structural remodeling, and endothelial-associated transcriptional programs were among the strongest contributors to human-aligned signatures. These data position the neurovascular unit as a driver of disease timing and phenotype, potentially delaying acute tauopathy while creating conditions for delayed crises that more closely resemble the staged evolution inferred from human CTE pathology ^7,11,33,38,39^.

In summary, this work addresses a major barrier in the field: the absence of a human experimental model that can capture the slow emergence of CTE-like cellular pathology after survivable repetitive mild injury. By establishing a multicellular human *in vitro* injury model with strong pathway-level convergence to human brain datasets, and by identifying endothelial/NVU and mitochondrial programs as key determinants of outcome, these findings reframe CTE-like progression as a state of maladaptive survival shaped by neurovascular-metabolic control. This system provides a tractable platform for defining early drivers, testing mechanism-based interventions that target mitochondrial function and neurovascular homeostasis, and developing biomarker-linked strategies to interrupt the transition from repetitive injury to chronic neurodegeneration ^7,36,40,49^.

## Limitations and Future Directions

This study has several limitations. It relies on a limited number of donor lines, and greater genetic diversity, including common susceptibility alleles such as APOE and MAPT, will be needed to capture patient variability. The current eight-week window revealed early tauopathy and metabolic remodeling but does not yet demonstrate frank neurodegeneration; extending cultures to six months or longer will be critical to assess neuronal loss and late-stage pathology. Additional complexity, such as incorporating oligodendrocytes and more physiologic injury paradigms, will further strengthen relevance. Finally, higher-resolution methods such as single-cell transcriptomics and functional assays are needed to define how specific cell types initiate and sustain disease. Importantly, the ability to dissect these mechanisms at the level of individual cell types creates opportunities for targeted therapies, such as stimulating mitochondrial resilience in the most vulnerable populations.

## Materials and Methods

### 3D silk scaffold preparation

Silk fibroin scaffolds were fabricated from *Bombyx mori* cocoons using established methods ^16,17,19,21,50^. Cocoons were boiled for 30 minutes in sodium carbonate to remove sericin, after which the isolated fibroin fibers were rinsed and dried overnight. Dried fibers were dissolved in 9.3 M lithium bromide and dialyzed against deionized water for three days to remove residual salt. The resulting silk fibroin solution was adjusted to 6% (w/v), cast into 6-cm diameter dishes, and combined with sodium chloride particles (300-425 μm) to generate porosity. After two days of incubation, the mixture was heated at 60°C for one hour to induce β-sheet formation. Scaffolds were then removed and dialyzed for an additional two days to leach out the salt, yielding three-dimensional porous sponges.

Cylindrical scaffolds were generated using biopsy punches (McMaster-Carr) to produce constructs with a 6-mm outer diameter and 2-mm central lumen, and trimmed to a height of 1.5 mm. Scaffolds were sterilized by autoclaving for 20 minutes and stored at 4°C for up to one week before use.

### Human brain cell culture preparation

#### Mouse Embryonic Fibroblasts (MEFs)

Mouse embryonic fibroblasts (MEFs) were cultured as feeder layers for human induced neural stem cells (hiNSCs) according to ATCC guidelines. MEFs (passages 2–4) were maintained in DMEM (1× + GlutaMAX; Thermo Fisher, 10569010) supplemented with 10% fetal bovine serum (FBS; Gibco, 10438-026) and 1% antibiotic–antimycotic (Thermo Fisher, 15240062) at 37°C and 5% CO_2_, with media changes every 3 days. Prior to seeding, 15-cm dishes were coated with 0.1% gelatin for 1 hour at room temperature.

For expansion, cells were detached using 0.25% Trypsin-EDTA for 3 minutes at 37°C, neutralized with MEF medium, and centrifuged at 1000 rpm for 5 minutes. At 90–100% confluency, MEFs were mitotically inactivated with mitomycin C (10 μg/mL; Sigma, M4287) for 3 hours at 37°C, washed three times with PBS, and either used immediately for hiNSC seeding or maintained in MEF medium for up to 7 days.

#### Human-Induced Neural Stem Cells [hiNSCs]

Human induced neural stem cells (hiNSCs) were generated from dermis-derived neonatal foreskin fibroblasts ^51^. Cells were maintained in KnockOut DMEM supplemented with 20% KnockOut Serum Replacement, 1% GlutaMAX, 1% antibiotic–antimycotic, 0.2% β-mercaptoethanol (Invitrogen), and basic fibroblast growth factor (bFGF; 10 μg/mL stock, added fresh to a final concentration of 40 ng/mL). Media was refreshed every 2 days.

hiNSCs were initially seeded onto mitotically inactivated MEF feeder layers. For feeder-free expansion, cells were passaged onto 15-cm dishes coated with Matrigel (1:20 dilution) following detachment with TrypLE for 1 minute at 37°C and replated in hiNSC medium.

#### Human Primary Astrocytes

Human primary astrocytes (Sciencell Research Laboratories, 1800) were maintained in astrocyte growth medium containing fetal bovine serum, astrocyte growth supplement, and antibiotic–antimycotic solution at 37°C and 5% CO_2_, with media changes every 3–4 days. For seeding, 15-cm dishes were coated with poly-L-lysine (2.5 μL PLL in 15 mL sterile water; Sigma, P4832) for 1 hour at room temperature. Cells were expanded at 70–80% confluency by detachment with 0.25% Trypsin-EDTA for 3 minutes at 37°C, neutralized with growth medium, and centrifuged at 1000 rpm for 5 minutes.

#### HMC3 Microglia

HMC3 human microglial cells (ATCC, CRL-3304) were cultured in Eagle’s Minimum Essential Medium (EMEM; Thermo Fisher, 50-188-268FP) supplemented with 10% FBS (Gibco, 10438-026) and 1% antibiotic–antimycotic. Cells were maintained at 37°C and 5% CO_2_, with media changes every 3–4 days. Cells were passaged at 70–80% confluency using 0.25% Trypsin-EDTA for 3 minutes, followed by neutralization, centrifugation at 1000 rpm for 5 minutes, and replating.

#### Endothelial cells

Human brain microvascular endothelial cells (HBMVECs; cAP-0002) were cultured in endothelial growth medium (EGM; cAP-02) containing 10% serum and growth supplements at 37°C and 5% CO_2_, with media changes every 3–4 days. Prior to seeding, 15-cm dishes were coated with Quick Coating Solution (cAP-01) for 5 minutes. Cells were expanded at 70–80% confluency by detachment with 0.25% Trypsin-EDTA for 3 minutes at 37°C, neutralized with EGM, and centrifuged at 1000 rpm for 5 minutes.

#### Pericytes

Human brain vascular pericytes (Sciencell Research Laboratories, 1200) were maintained in pericyte growth medium containing fetal bovine serum, pericyte growth supplement, and antibiotic–antimycotic solution at 37°C and 5% CO_2_, with media changes every 3–4 days. Prior to seeding, 15-cm dishes were coated with poly-L-lysine (2.5 μL in 15 mL sterile water) for 1 hour at room temperature. Cells were expanded at 70–80% confluency using 0.25% Trypsin-EDTA for 3 minutes at 37°C, followed by neutralization, centrifugation at 1000 rpm for 5 minutes, and replating.

### Mycoplasma testing

All cell lines used in this project were either checked for mycoplasma before starting or were directly purchased from established vendors that provided certification for Mycoplasma testing.

### 3D model fabrication

The 3D multicellular engineered brain microvascular tissue model was adapted from our previously published work ^16,17^. Silk scaffolds were coated with poly-L-lysine (0.1 mg/mL, overnight at 37°C) followed by laminin (50 μg/mL, overnight at 4°C), washed with PBS, and conditioned in neural basal (NB; PENTA cultures). Neurons (50 million cells/mL), astrocytes (12.5 million cells/mL), and microglia (2.5 million cells/ml) were seeded onto dried scaffolds (40 μL per scaffold) and incubated for 30 minutes at 37°C to promote attachment, after which culture medium was added.

The following day, scaffolds were transferred to fresh plates and overlaid with a collagen type I hydrogel (100 μL, 3 mg/mL, pH 7.0) containing endothelial cells (10 million cells/mL) and pericytes (3 million cells/mL) and 0.5% matrigel. After collagen polymerization, constructs were cultured in NB or endothelial medium and transferred to 48-well plates the next day. Neurobasal medium was supplemented with astrocyte growth factors. Media was refreshed every four days until constructs were fixed or frozen at designated time points (2, 4, or 10 weeks) or subjected to controlled cortical impact 5–6 weeks post-seeding.

### Mechanical contusion injury model

Scaffolds were mechanically injured to simulate moderate TBI through the use of a controlled cortical impactor (CCI) device. Scaffolds were placed on sterile weigh boats within a biosafety cabinet and subjected to mechanical injury using a pneumatic CCI device in a manner similar to our group’s previously published model ^16,17,19^.

For the injury paradigm in the first experiment, which aimed to determine the threshold that permits survivable trauma for long-term modeling, impactor tips with diameters of 1 mm and 3 mm were employed at velocities of 1 m/s, 3 m/s, and 6 m/s, with an impact depth of 0.6 mm and a dwell time of 200 milliseconds. Furthermore, a series of injuries caused by 1 to 3 impacts were analyzed utilizing impactor tips with diameters of 1 mm and 3 mm at a velocity of 1 m/s. Consecutive injuries were administered when multiple injuries occurred.

In the second experiment, the injury paradigm aimed to assess how varying intervals between three mild repetitive injuries impact the 3D brain model. For the mild injury paradigm, a 1 mm diameter impactor tip was employed at a velocity of 1 m/s, with a total of three injuries at various intervals, including consecutive injuries, 1-day, 3-day, 7-day, and 14-day intervals. For further experiments on the mild repetitive injury condition, three mild injuries (1 mm tip size and 1 m/s speed) with a 14-day interval were used.

Injuries were administered 6 weeks after scaffold seeding, with each injured sample receiving an injury based on the injury condition. Sham controls were exposed to identical environmental conditions, including air exposure on the weigh boat, but did not undergo mechanical impact. Sample collection timepoints were defined relative to the injury event.

### Western blot

Protein expression was quantified by Western blotting. Whole tissue scaffolds stored at −80°C were lysed in 1× RIPA buffer and subjected to brief sonication (20% amplitude, 20 pulses; 1 s on/1 s off) to facilitate protein extraction. Lysates were combined with sample buffer (Laemmli buffer supplemented with β-mercaptoethanol at a 9:1 ratio) at a 3:1 lysate-to-buffer ratio. Mitochondrial and total protein fractions obtained following Seahorse assay processing were prepared in parallel. All samples were denatured at 95°C for 5 minutes prior to electrophoresis.

Proteins (15 μL lysate plus 5 μL sample buffer) were separated by SDS-PAGE at 150 V until dye front migration was complete (∼50 minutes). Precision Plus Protein™ All Blue Standards (Bio-Rad) were used for molecular weight reference. Gels were imaged using stain-free detection prior to transfer. When sample numbers exceeded single-gel capacity, samples were distributed across multiple gels and sectioned at consistent molecular weight positions (75 and 20 kDa) to enable uniform transfer.

Proteins were transferred to PVDF membranes using a semi-dry transfer system (2.5 V, 1.6 A) for 4 minutes for total protein samples or 3 minutes for mitochondrial fractions. Membranes were imaged post-transfer using stain-free detection to quantify total protein for normalization. Membranes were probed sequentially with primary antibodies (typically 1:1000; 1:3000 for high-abundance targets) overnight at 4°C, followed by incubation with HRP-conjugated secondary antibodies (1:10,000) for 1 hour. Washes were performed in TBST between all incubation steps. Signal detection was carried out using enhanced chemiluminescence, with membranes exposed for 5 minutes prior to imaging. For multi-target analysis, membranes were stripped and reprobed as needed.

Densitometric quantification was performed by normalizing all targets to total protein per lane. Phosphorylation ratios were calculated as pTau/Tau and subsequently normalized to total protein. All other protein levels are reported relative to total protein content.

Western blot Primary Antibody: TOMM20 Rb (Invitrogen REF MA5-24859), pTau (Ser202 & Thr205) Ms (Invitrogen REF MN1020), pTau Rb (THR231, Cell Signaling 71429S), pTau Rb (PHF-1, Invitrogen PIPA5101061), SYN1 Rb (Abcam AB254349), MAP2 Rb (NC0165384), TDP43 Rb (Abcam ab109535), APP Rb (Invitrogen REF PA5-17829), Tau Ms (Cell Signaling 4019S), Tuj1 Rb (Abcam ab18207) . Western Blot Secondary Antibody: Anti-rabbit IgG, HRP-linked (Cell Signaling 7074), Anti-mouse IgG, HRP-linked (Cell Signaling 7076)

### Immunofluorescence and confocal imaging

Cellular morphology and protein localization were assessed by immunofluorescence and confocal microscopy. Three-dimensional cell-seeded scaffolds were fixed in 4% paraformaldehyde with 4% sucrose in phosphate-buffered saline (PBS) for 1 hour at room temperature, washed in PBS, and permeabilized for 1 hour in PBS containing 0.2% Triton X-100 and 4% goat serum. Primary antibodies were applied overnight at 4°C in permeabilization buffer. Samples were washed with PBS and incubated with fluorophore-conjugated secondary antibodies for 1 hour at room temperature, followed by nuclear staining with DAPI for 5 minutes. After final washes, fluorescent image stacks were acquired using a Zeiss confocal microscope.

Regions of interest were selected blinded to the quantification channel (pTAU, Abeta, or TDP43). For each sample, 50 z-stacks were collected at 10× zoom (1024 × 1024 pixels; 3181.98 × 3181.98 μm field of view) with a step size of 0.34 μm. Images are presented as maximum-intensity projections of combined channels. Image adjustments were applied uniformly for figure presentation only and were not used for quantitative analysis.

### Neuronal network analysis

Neuronal network density was quantified from Tuj1-labeled z-stacks using a custom MATLAB pipeline previously described by the Liaudanskaya laboratory ^19^. Background signal from the silk scaffold and neuronal cell bodies was removed using Otsu thresholding followed by eccentricity filtering (≥0.90) to retain elongated neurite structures. Neurite voxels were grouped using 18-point connectivity (voxel size: 0.258 × 0.258 × 0.422 μm), and neurite length was calculated as the longest principal axis of the best-fit ellipsoid for each connected object. Network prevalence was defined as the proportion of voxels included in neurite analysis relative to the total voxel count per stack.

### Vascular network analysis

Vascular-like network architecture was quantified using AngioTool, an NIH-developed open-source software for angiogenesis analysis ^17^. Analyses were performed on maximum-intensity projections of the GFP channel with vessels segmented against a dark background. Images were processed using a multiscale Hessian-based enhancement filter to detect tubular structures, followed by skeletonization. Quantified metrics included total and average vessel length, vessel area, junction number, vascular density, lacunarity, and branching index (branch points per unit area). Lacunarity was calculated using a fast box-counting algorithm. Identical scale and segmentation parameters were applied across all experimental groups.

Primary antibodies used were Tuj1/βIII-tubulin (Chicken; Fisher NB1001612), pTau (Ser202 & Thr205) Ms (Invitrogen REF MN1020), TDP43 Rb (Abcam ab109535), and Amyloid Fibril (rabbit; Abcam ab201062). Secondary antibodies included goat anti-chicken IgY (H+L) (Invitrogen, A32933), goat anti-mouse IgG (H+L) (Invitrogen, A32728), and goat anti-rabbit IgG (H+L) (Invitrogen, A32733).

### Neurodegeneration panel

Media samples were collected from scaffold culture experiments and stored at -80°C until processing. Prior to assay, frozen samples were thawed on ice and centrifuged at 1,400 rpm for 10 minutes at 4 °C to remove cell debris. The supernatant was transferred to new Eppendorf tubes for the analysis.

Samples were sent to the research flow cytometry facility at Cincinnati Children’s Hospital Medical Center. Neurodegenerative protein levels were measured using the Luminex platform with the ProcartaPlex™ Human Neurodegeneration Panel 1, 9plex (ThermoFisher, EPX090-15836-901).

### Seahorse assay

Mitochondrial bioenergetic function was evaluated using the Seahorse XFe96 Analyzer and the Real-Time ATP Rate Assay Kit (Agilent Technologies, 103792-100) following the manufacturer’s protocol with minor modifications optimized for 3D silk scaffold-derived samples.

#### Sensor and plate preparation

Approximately 12–18 hours prior to the assay, Seahorse sensor cartridges were hydrated by adding 200 µL of sterile water to each well of the utility plate and incubated overnight in a 37°C, non-CO_2_ incubator. On the day of the assay, water was replaced with 200 µL of XF Calibrant, and the sensors were equilibrated for at least 2–4 hours at 37°C.

#### Assay medium and drug preparation

Assay medium consisted of phenol red–free DMEM/F12 supplemented with 10 mM glucose, 1 mM pyruvate, and 2 mM glutamine (pH 7.4). On the day of the assay, stock solutions were freshly prepared to achieve final working concentrations of 15 µM oligomycin and 5 µM rotenone/antimycin A after injection. The drugs were loaded into the sensor cartridge (20 µL of oligomycin in Port A; 22 µL of rotenone/antimycin A in Port B).

#### Mitochondrial isolation

Mitochondria were isolated from 3D silk scaffold cultures at the specified time point after treatment using the Mitochondria Isolation Kit (ThermoFisher) with optimized handling for silk-based tissues. For intracellular mitochondria, scaffolds were transferred to 2 mL microtubes containing 800 µL of Isolation Reagent A and disrupted mechanically. After brief vortexing and incubation on ice, 10 µL of Reagent B was added, followed by further incubation and addition of 800 µL of Reagent C. The scaffolds were removed, and the suspension was centrifuged at 1,000 × g for 10 minutes at 4°C to remove debris. The resulting supernatant was centrifuged at 12,000 × g for 25 minutes at 4°C to pellet mitochondria. Extracellular mitochondria were isolated from conditioned media using the same differential centrifugation procedure.

#### Seahorse assay setup and data collection

Mitochondrial pellets were resuspended in 180 µL of Seahorse assay medium and loaded into designated wells of a Seahorse XF cell culture microplate. Blank wells received assay medium only. Following sensor calibration, the loaded cell culture plate was inserted into the Seahorse XFe96 Analyzer and run using the “Real-Time ATP Rate Assay” template. Oxygen Consumption Rate (OCR) and Extracellular Acidification Rate (ECAR) were measured in real time to quantify ATP production from oxidative phosphorylation and glycolysis, respectively.

#### Post-assay sample collection

Upon completion, 20 µL of the mitochondria-rich media from each well was stored at −20°C for mitochondrial DNA (mtDNA) quantification. The remaining 160 µL was centrifuged at 12,000 × g for 20 minutes, and the mitochondrial pellet was reconstituted in 60 µL of RIPA buffer (supplemented with protease and phosphatase inhibitors) for storage and subsequent protein analysis.

### RNA isolation protocol

Scaffolds were collected and stored at −80°C. For RNA isolation, samples were thawed on ice and incubated in 600 μL of RNeasy lysis buffer (Qiagen, 74106). During lysis, scaffolds were mechanically disrupted using fine scissors and briefly sonicated to facilitate tissue dissociation (20% amplitude, 5 s total; 1 s on/1 s off). Lysates were incubated on ice for an additional 20 minutes and then passed through QIAshredder Mini Spin columns (Qiagen) to remove residual silk material, followed by centrifugation at 15,000 rpm for 2 minutes.

An equal volume of 70% ethanol was added to the cleared lysate and mixed thoroughly. Samples were loaded onto RNeasy Mini Spin columns (Qiagen) in two sequential centrifugation steps (10,000 rpm, 1 minute each), discarding the flow-through after each spin. Columns were washed with 350 μL of RNeasy wash buffer and incubated for 5 minutes prior to centrifugation. On-column DNase digestion was performed using TURBO DNase according to the manufacturer’s instructions (Invitrogen, P5537155). Following DNase treatment, columns underwent additional wash steps and a final dry spin (10,000 × g, 1.5 minutes). RNA was eluted in 24 μL of RNase-free water (Fisher Scientific) with a 10-minute incubation, followed by two sequential centrifugation steps (10,000 rpm, 1 minute each). RNA concentration was quantified using a Tecan Infinite M Plex NanoQuant Plate™, and samples were stored at −80°C prior to sequencing.

RNA sequencing was performed by Azenta Life Sciences. FASTQ files were aligned using HISAT2, and gene-level counts were generated with featureCounts. Ensembl gene identifiers were converted to HGNC symbols using biomaRt in R. Count normalization and differential expression analyses were performed using DESeq2. Gene set enrichment analysis (GSEA) was conducted to assess differences between CCI and SHAM conditions within PENTA and NAM cultures using MitoCarta 3.0 pathway annotations. Log_2_ fold-change values (CCI:SHAM) were used for comparative analyses between PENTA and NAM models. Data visualization was performed using ggplot2, including generation of lollipop plots.

### RNA sequencing analysis and cross-platform transcriptomic integration

#### Bulk RNA-seq preprocessing and experimental grouping

To compare injury-responsive transcriptional programs across *in vitro* models and human chronic traumatic encephalopathy (CTE), bulk RNA-sequencing data were analyzed from two human 3D culture platforms, NAM and PENTA, collected at acute (24 h) and chronic (8 weeks) time points following controlled cortical impact (CCI), together with published human CTE transcriptomic datasets ^9,25^. Gene-level count matrices for NAM and PENTA cultures were provided as comma-separated value files with genes in rows and samples in columns. No samples were excluded from analysis.

Sample metadata, including injury status (CCI vs sham) and time point, were inferred directly from sample identifiers using predefined, case-insensitive regular expression matching. This yielded four experimental groups per model: sham-24 h, CCI-24 h, sham-8 weeks, and CCI-8 weeks. All statistical comparisons were performed using time-matched sham controls.

Gene identifiers were harmonized across datasets by trimming whitespace and converting all identifiers to uppercase official gene symbols. For exploratory analyses, raw counts were transformed as log2(count + 1) to stabilize variance.

#### Gene-level injury screening

Within each *in vitro* model and time point, injury-associated expression changes were quantified as the difference in mean log2-transformed expression between CCI and sham samples. Statistical significance at the gene level was assessed using two-sided Welch’s t-tests, which do not assume equal variance and are appropriate for modest and unequal sample sizes. Gene-level p-values were used for descriptive summaries, result export, and annotation, but were not used as the primary basis for biological interpretation.

### Human CTE module-based analysis

#### Definition and projection of human CTE co-expression modules

Human CTE-associated co-expression modules and corresponding differential expression values were obtained from published postmortem brain datasets ^9,25^. Module definitions were treated as fixed gene sets and were not re-derived. Only genes present in both the human and *in vitro* datasets were retained. Modules containing fewer than 10 overlapping genes were excluded.

For each remaining module, coordinated transcriptional activity (“module program activity”) was quantified in NAM and PENTA datasets using principal component analysis (PCA). Log2(count + 1) expression values for module genes were z-scored across genes to ensure equal weighting. PCA was performed on the transposed matrix (samples × genes), and the first principal component (PC1) score was used as the module activity score. PC1 signs were adjusted so that the mean score across samples was positive, enabling consistent interpretation of directionality.

#### Module-level injury effects and statistical testing

Within each model and time point, injury effects were quantified as the difference in mean PC1 score between CCI and sham samples. Statistical significance was assessed using two-sided Welch’s t-tests on PC1 scores. Effect sizes were reported as raw differences (CCI − sham), and variability was summarized using the standard error of the mean.

To formally assess temporal divergence of injury responses, difference-in-differences analyses were performed using linear models of the form:

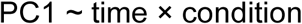

where time (24 h vs 8 weeks) and condition (CCI vs sham) were treated as categorical variables.

The interaction term tested whether injury effects differed between acute and chronic time points. Interaction p-values were used to identify modules with temporally distinct injury responses. These models were fit using unnormalized PC1 scores.

For visualization only, PC1 scores were mean-centered to time-matched sham samples, and sham samples from both time points were pooled. This normalization was not used for statistical testing.

#### Human- in vitro module concordance

Human CTE module effects were summarized by averaging reported human differential expression values across genes within each module. Where sample-level human data were available, module scores were computed per sample and compared between CTE and control groups using Welch’s t-tests. Module-level effect sizes from human, NAM, and PENTA datasets were jointly visualized using heatmaps, optionally sorted by acute in vitro injury effects to emphasize early responses.

#### Cross-platform matching with human single-nucleus RNA-seq

To identify culture-specific injury responses, differential expression screening was performed separately for NAM and PENTA cultures at 24 h and 8 weeks. Raw counts were normalized to counts per million (CPM), filtered to retain genes with CPM > 1 in at least two samples, and log-transformed as log2(CPM + 1). Differential expression was assessed using two-sided Welch’s t-tests with Benjamini–Hochberg false discovery rate (FDR) correction. Genes were considered injury-responsive if FDR < 0.05 and |log2FC| > 0.25.

Culture-specific signatures were defined as:

- PENTA-only: significant in PENTA but not NAM at the same time point
- NAM-only: significant in NAM but not PENTA at the same time point

#### Integration with human snRNA-seq datasets

Human single-nucleus RNA-seq differential expression results were obtained from published data ^9^. For each cell type, genes with FDR < 0.05 in RHI vs control or CTE vs control comparisons were retained. Oligodendrocytes and T cells were excluded a priori.

Cross-platform overlap was restricted to directionally concordant genes, defined as genes that were (i) culture-specific in vitro, (ii) significant in human data, and (iii) changed in the same direction in both datasets. Overlaps were computed separately for RHI and CTE, and the union was used for visualization.

Enrichment of human cell-type DEGs within culture-specific signatures was assessed using one-sided Fisher’s exact tests with Benjamini–Hochberg correction. Directional agreement was quantified using Spearman correlation between in vitro and human log2FC values when at least five intersecting genes were present.

#### Mitochondrial pathway analysis

To specifically evaluate mitochondrial involvement in injury responses, mitochondrial gene and pathway analyses were performed using curated mitochondrial annotations from MitoCarta 3.0. For each *in vitro* model, time point, and injury condition, mitochondrial pathway activity was assessed using both gene-level and module-level approaches.

At the gene level, injury-associated log2FC values (CCI vs time-matched sham) were extracted for all MitoCarta genes. For pathway-level analyses, mitochondrial genes were grouped into functional categories defined by MitoCarta 3.0 (e.g., oxidative phosphorylation, mitochondrial translation, mitochondrial dynamics, and metabolic pathways). For each pathway, an aggregate pathway score was computed per sample as the mean log2-transformed expression across all genes assigned to that pathway.

Pathway scores were compared between CCI and sham samples using two-sided Welch’s t-tests at each time point. Directionality of regulation was defined relative to time-matched sham controls. For visualization, pathway scores were displayed as absolute values to illustrate baseline maturation trajectories and injury-induced deviations. Upward bars indicate pathways enriched among genes upregulated relative to sham, whereas downward bars indicate enrichment among downregulated genes.

To assess concordance with human disease, mitochondrial pathway gene sets were intersected with directionally concordant human CTE and RHI DEGs using the criteria described above. Over-representation analyses were performed using one-sided Fisher’s exact tests with FDR correction. Pathway-level summaries and gene-count–based enrichments were presented as complementary measures of mitochondrial involvement.

Protein-protein interaction and pathway enrichment analyses for mitochondrial gene sets were performed using STRING (v11+ API), with Homo sapiens as the reference organism and a minimum interaction score of 0.4. Network hubs were identified using node degree and betweenness centrality metrics.

#### Statistical considerations and software

All RNA sequencing statistical analyses were performed using Python and R. Bulk RNA-seq preprocessing, normalization, differential expression screening, module projection, and cross-platform matching were implemented in Python using custom analysis pipelines. Pathway enrichment analyses and pathway-level visualizations were performed in R. Nominal p values are reported unless otherwise indicated, with multiple-testing correction applied using the Benjamini-Hochberg procedure where appropriate. All figures were generated using standardized visualization parameters to facilitate direct comparison across experimental conditions.

### Statistical analysis

Statistical analyses were performed using GraphPad Prism 10. Two-tailed t tests were used for comparisons between two experimental groups. One-way analysis of variance (ANOVA) was used for comparisons among multiple groups within a single time point or across multiple time points within a single group, followed by Tukey’s post hoc test for pairwise comparisons and comparison to control conditions. A p value ≤ 0.05 was considered statistically significant.

All experiments were independently repeated at least three times to minimize *in vitro* variability. Because each scaffold was cultured independently, individual scaffolds were treated as independent samples. Within each experimental replicate, data were normalized to the mean of the corresponding negative control group before combining replicates for statistical analysis. For example, for western blot signals (Tuj1 normalized to total protein per lane) were further normalized to the mean of the Sham group within each experiment. Data are presented as mean ± standard error of the mean (SEM) unless otherwise specified.

## Ethics Declaration

### Ethics

This work did not include any experiments with animals or the participation of human subjects.

## Authors contribution

VL conceived the project. SJ and VL designed and interpreted the experiments and wrote the manuscript. SJ and DSH performed scaffold seeding and sample maintenance (media changes). SJ and DSH performed all *in vitro* experiments, injuries. SJ, DSH, and JT performed mitochondria bioenergetic function assessment. SJ performed confocal imaging and network analysis. SJ, DSH, JT, CCA, HR, CMA, and MDT conducted Western Blots. SJ performed a multiplex panel. SJ, DSH, JT, CCA, HR, CMA, and MDT prepared silk materials. SBSK and VL performed RNA sequencing analysis.

## Acknowledgements

Cincinnati Children’s Hospital Bioimaging and Analysis Facility, and Research Flow Cytometry Facility. Naomi Habib’s group from Hebrew University for initial assistance with RNA sequencing results.

