## Supplement for "Injury-induced CTE-like pathology emerges in a human multicellular *in vitro* brain model and reveals mitochondrial and neurovascular regulation"

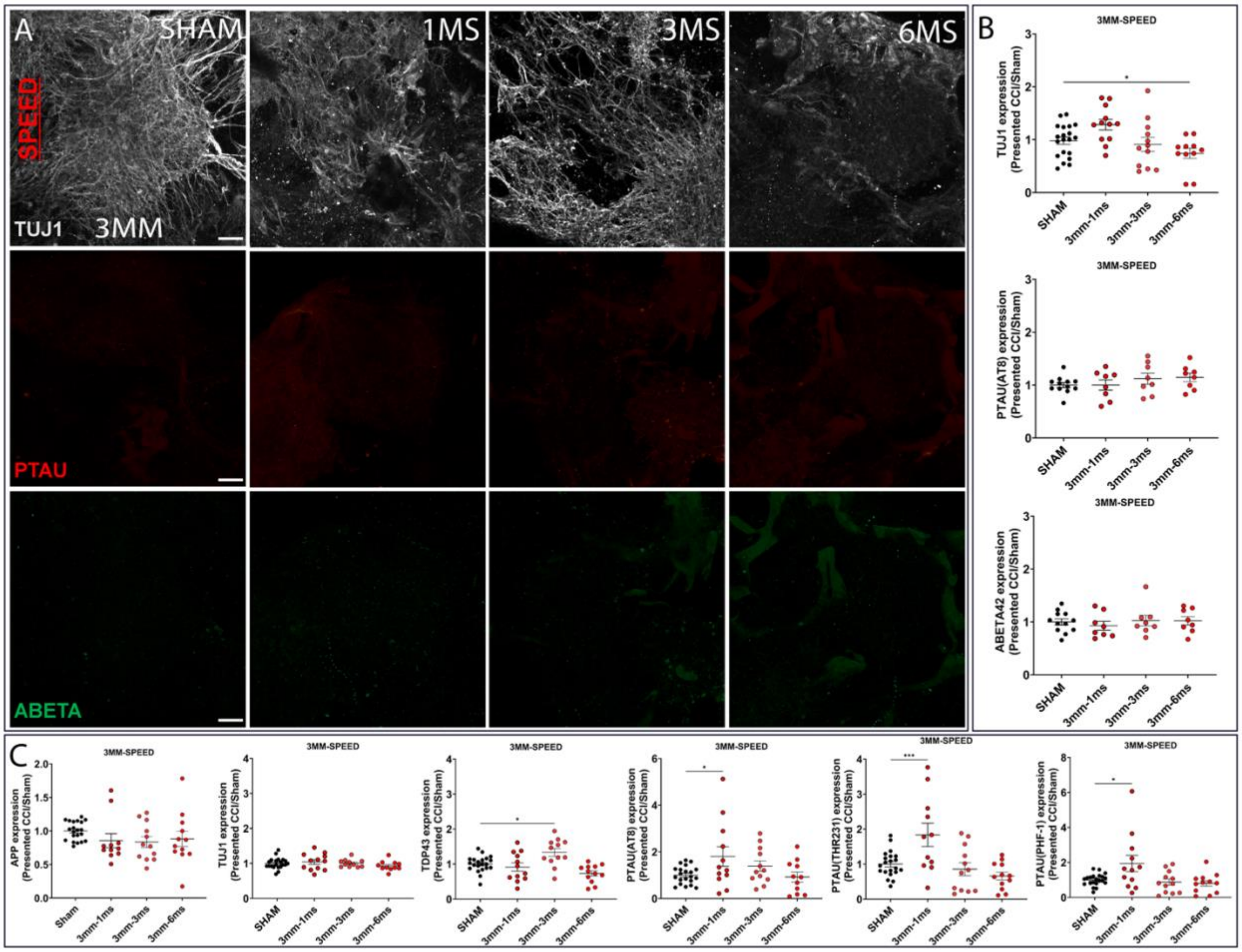


**Supplementary Figure 1.** Experiments were conducted to assess the effects of varying speeds (1m/s, 3m/s, and 6m/s) with a 3 mm tip size. **A** Representative image of NAM tri-culture 3 mm tip size with various speeds mentioned above. Immunofluorescent imaging of Tuj1 marker for neuronal networks; phosphorylated TAU(AT8) for degenerating neurons; and Ab42 for pathological proteinopathy. Scale bar: 100 µm. **B** Quantifications of the images from panel **A** (n = 6 – 10 scaffolds per condition). **C** Intracellular markers – Western Blot quantification of AD-related (APP, TDP-43), CTE-related (pTAU(AT8), pTAU(THR231), and pTAU(PHF-1)), and neuronal pan (Tuj1) markers (n = 9 – 12 scaffolds per condition) for speed. Data presented mean ± SEM of n=6-12 scaffolds per condition. *, *** indicates a significant difference with p<0.05, p<0.001, respectively. A one-way ANOVA (analysis of variance) test was used to determine the difference between the control and experimental groups. The ROUT outlier analysis method was used to exclude statistical outliers. Experiments were replicated at least three times, unless specified otherwise. Confocal images of pTau(AT8) and Ab42 were replicated two times.

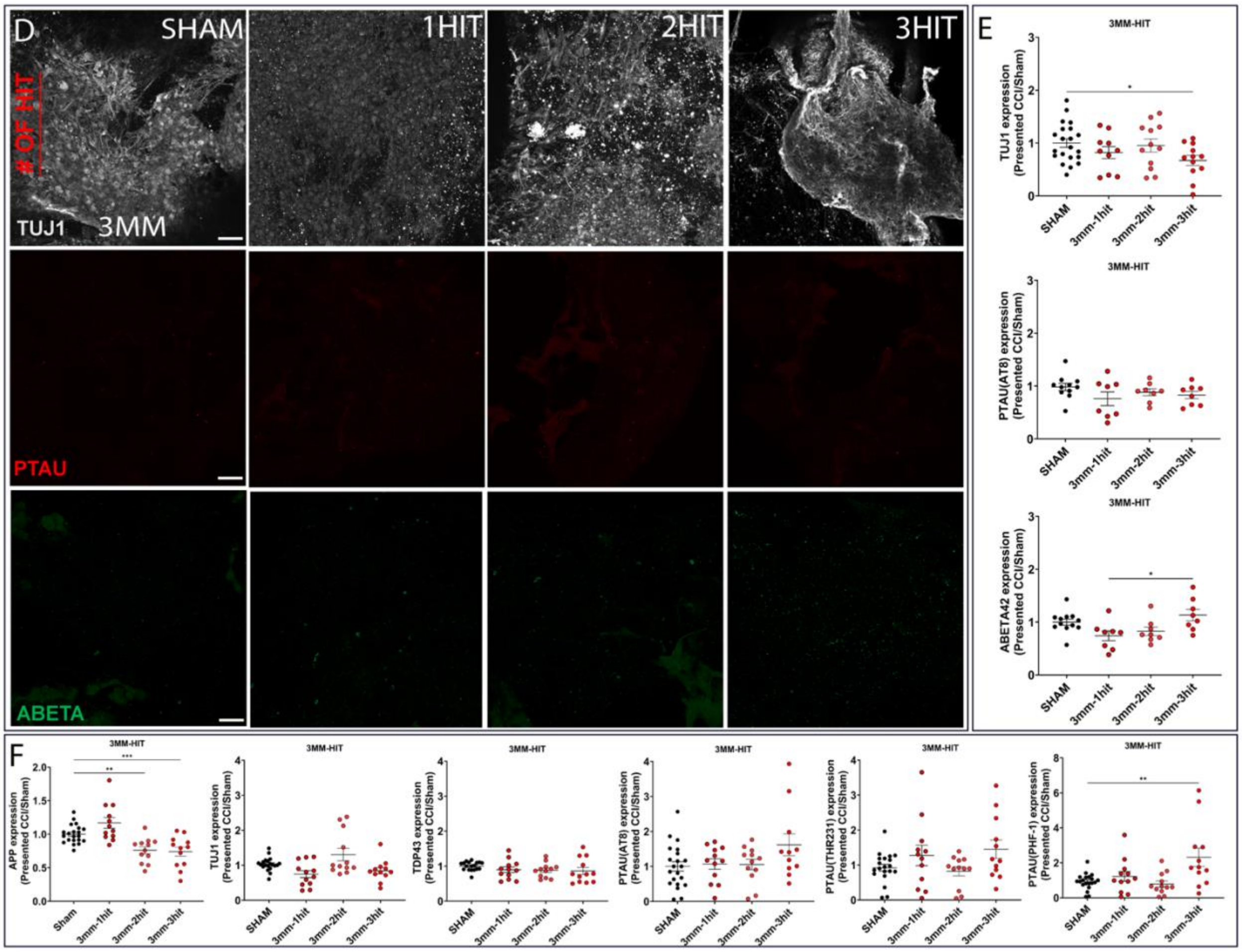


**Supplementary Figure 2.** Continuation of *Supplementary Figure 1* demonstrating a 3 mm tip size injury with different injury counts (1 hit, 2 hits, and 3 hits). **D** Representative image of NAM tri-culture 3 mm tip size with various injury counts mentioned above. Immunofluorescent imaging of Tuj1 marker for neuronal networks; phosphorylated TAU(AT8) for degenerating neurons; and Ab42 for pathological proteinopathy. Scale bar: 100 µm. **E** Quantifications of the images from panel **D** (n = 6 – 10 scaffolds per condition).

**F** Intracellular markers – Western Blot quantification of AD-related (APP, TDP-43), CTE-related (pTAU(AT8), pTAU(THR231), and pTAU(PHF-1)), and neuronal pan (Tuj1) markers (n = 9 – 12 scaffolds per condition) for speed. Data presented mean ± SEM of n=6-12 scaffolds per condition. ***,** **, *** indicates a significant difference with $p<0.05$, $p<0.01$, $p<0.001$, respectively. A one-way ANOVA (analysis of variance) test was used to determine the difference between the control and experimental groups. The ROUT outlier analysis method was used to exclude statistical outliers. Experiments were replicated at least three times, unless specified otherwise. Confocal images of pTau(AT8) and Ab42 were replicated two times.

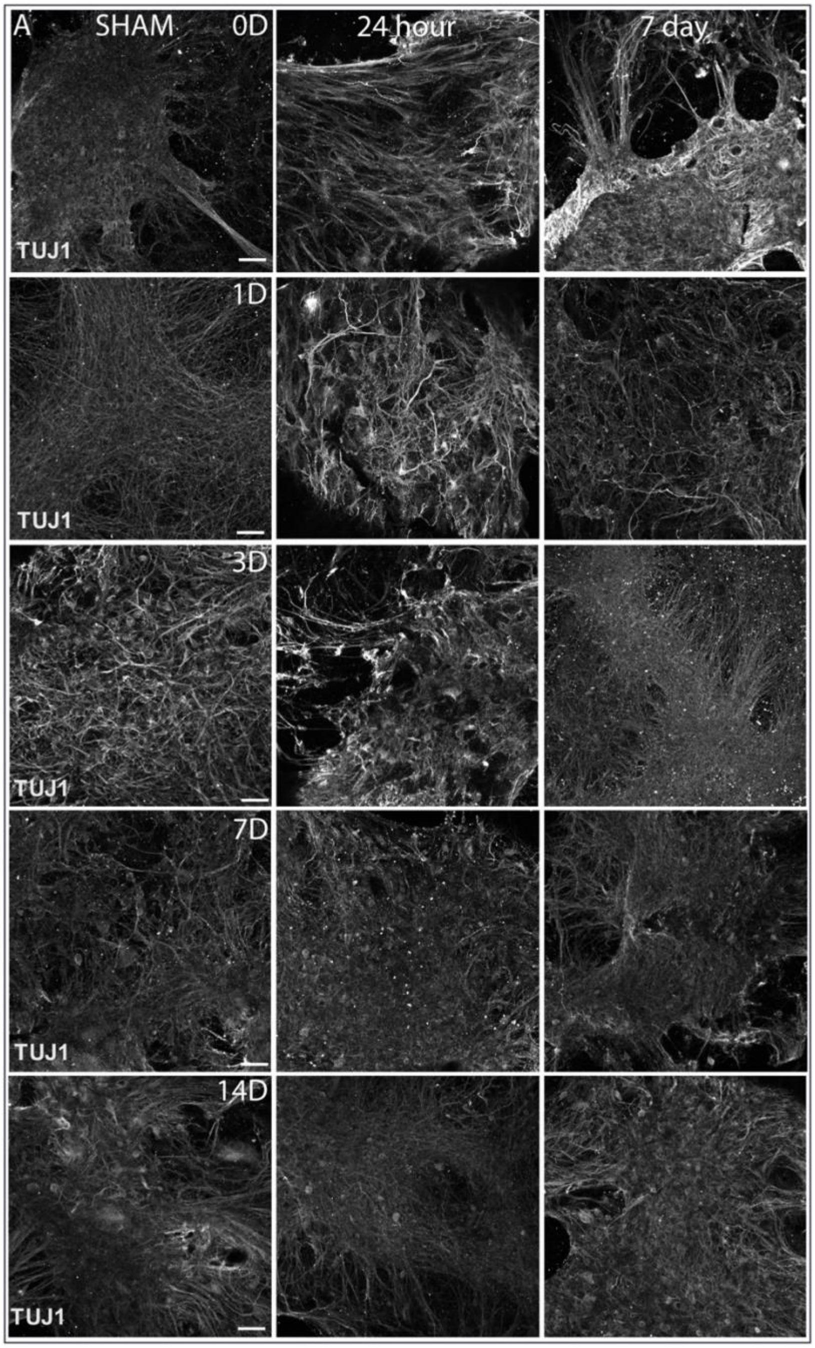


**Supplementary Figure 3.** Three mild repetitive injuries (1 mm-1 m/s) were applied to the NAM triculture model with various intervals (0-day, 1-day, 3-day, 7-day, and 14-day intervals). Representative Tuj1 (pan neuronal marker) staining of sham, 24 hours, and 7 days post-injury condition. Scale bar: 100 µm.

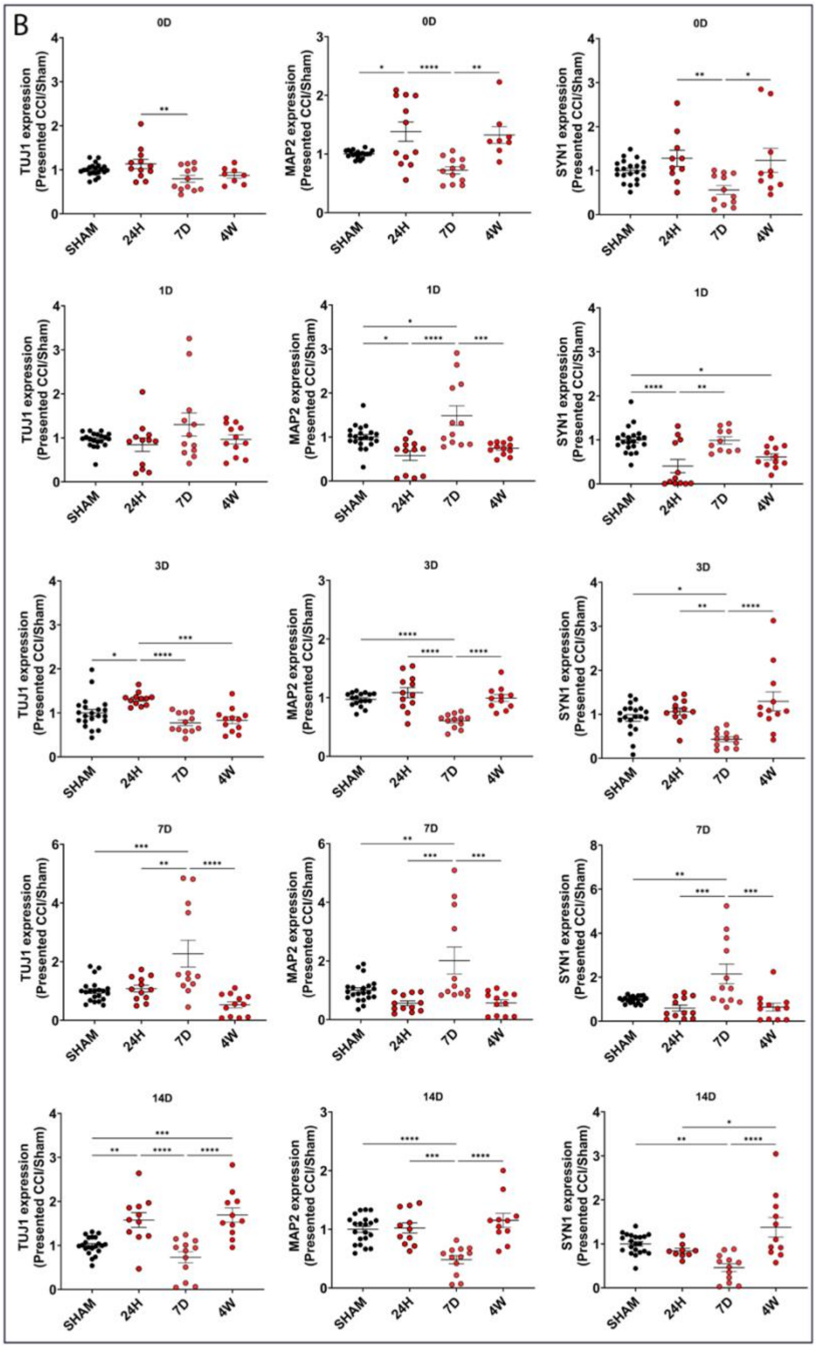


**Supplementary Figure 4.** Continuation of *Supplementary Figure 3*. **B** Intracellular markers – Western Blot quantification of Neuronal structure and Synaptic health (Tuj1, MAP2, and SYN1) markers.

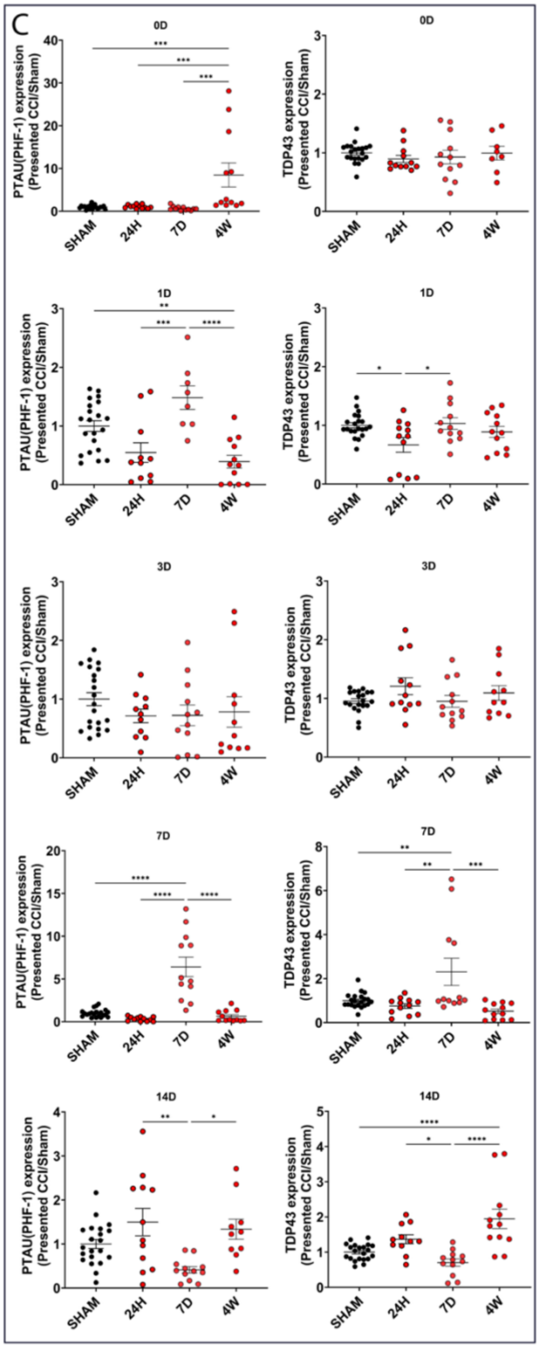


**Supplementary Figure 5.** Continuation of *Supplementary Figure 4*. **C** Intracellular markers – Western Blot quantification of Tau and non-Tau pathological (pTAU (PHF-1) and TDP-43) markers. Data presented mean ± SEM of n = 9 – 12 scaffolds per condition. ***,** **, ***, **** indicates a significant difference with p<0.05, p<0.01, p<0.001, p<0.0001, respectively. A one-way ANOVA (analysis of variance) test was used to determine the difference between the control and experimental groups. The ROUT outlier analysis method was used to exclude statistical outliers. Experiments were replicated at least three times.

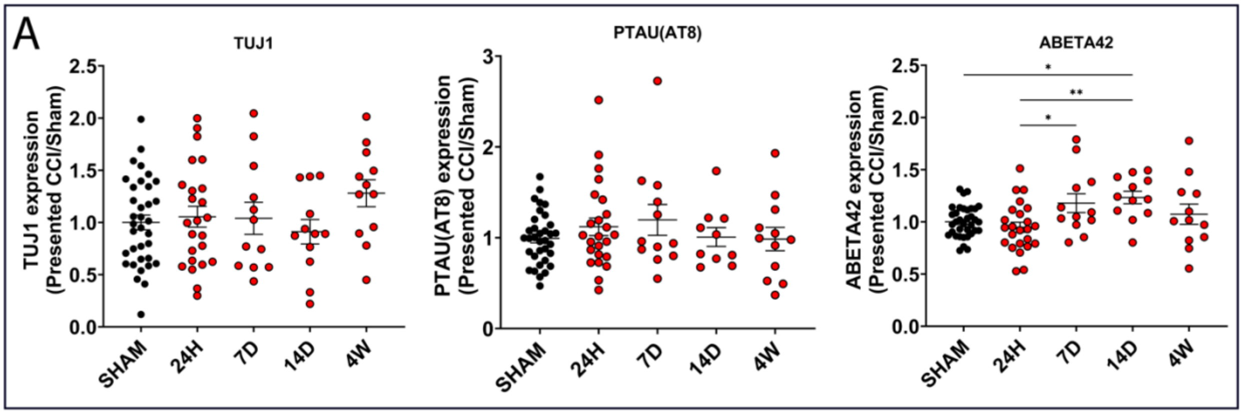


**Supplementary Figure 6.** Three mild repetitive injuries (1 mm-1 m/s) were applied to the NAM-triculture model with a 14-day interval, incorporating extended time points at 24 hours, 7 days, 14 days, and 4 weeks. **A** Quantifications of the immunofluorescent images of pan neuronal marker (Tuj1); phosphorylated TAU(AT8) for degenerating neurons, and Ab42 for pathological proteinopathy. Data presented mean ± SEM of n = 6 – 12 scaffolds per condition. ***,** **, indicates a significant difference with $p<0.05$, $p<0.01$, respectively. A one-way ANOVA (analysis of variance) test was used to determine the difference between the control and experimental groups. The ROUT outlier analysis method was used to exclude statistical outliers. Experiments were replicated at least three times, unless specified otherwise.

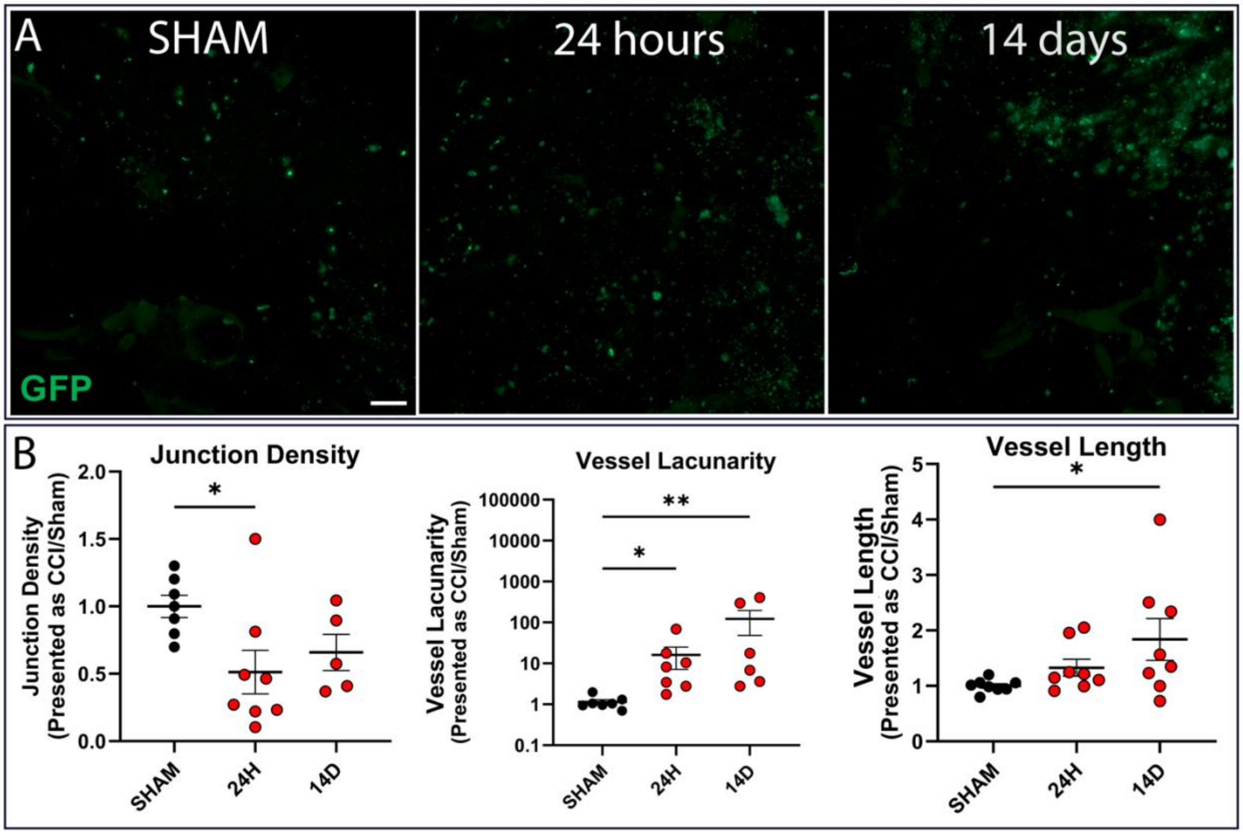


**Supplementary Figure 7.** Three mild repetitive injuries (1 mm-1 m/s) were applied to the PENTA culture model with a 14-day interval, incorporating extended time points at 24 hours and 14 days. **A** Confocal immunofluorescence image with hBMEC-produced GFP (488). Scale bar: 100 µm. **B** AngioTool quantification of vascular-like network lacunarity, junction density, and vessel length normalized to sham. Data presented mean ± SEM of n = 6 – 10 scaffolds per condition. ***,** **, indicates a significant difference with $p<0.05$, $p<0.01$, respectively. A one-way ANOVA (analysis of variance) test was used to determine the difference between the control and experimental groups. The ROUT outlier analysis method was used to exclude statistical outliers. Experiments were replicated two times.

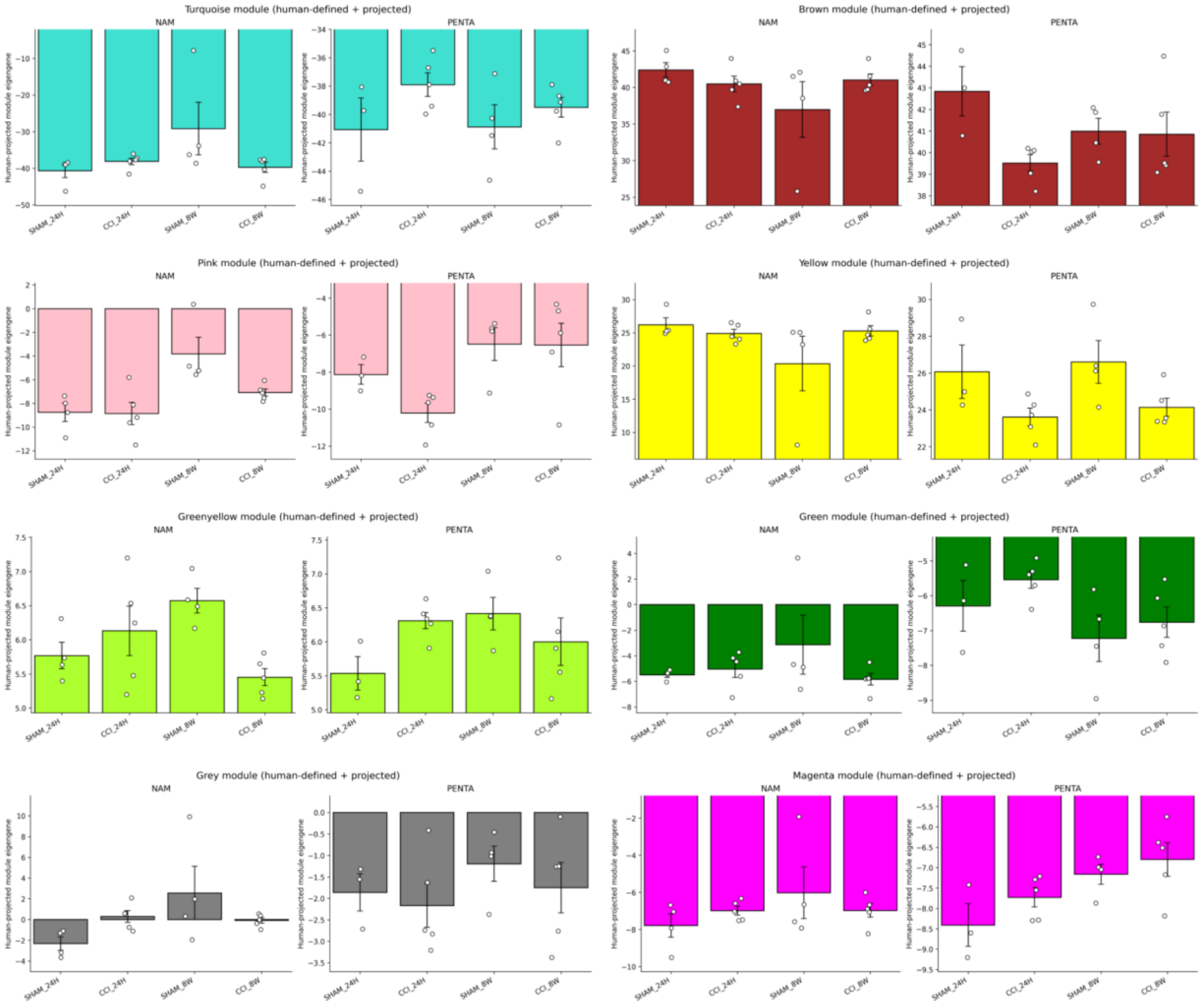


**Supplementary Figure 8.** Differential engagement of human CTE-associated gene modules in NAM and PENTA cultures following injury. Human CTE-associated gene modules were projected onto bulk transcriptomic profiles from NAM and PENTA cultures following sham or repetitive mild injury. Bar plots show module eigengene expression for representative human-defined modules (Turquoise, Brown, Pink, Yellow, Greenyellow, Green, Grey, and Magenta), displayed separately for NAM and PENTA systems. For each module, expression is shown for sham and post-injury conditions at 24 hours and 8 weeks, normalized to the corresponding sham control. Individual data points represent biological replicates; bars indicate mean ± SEM.

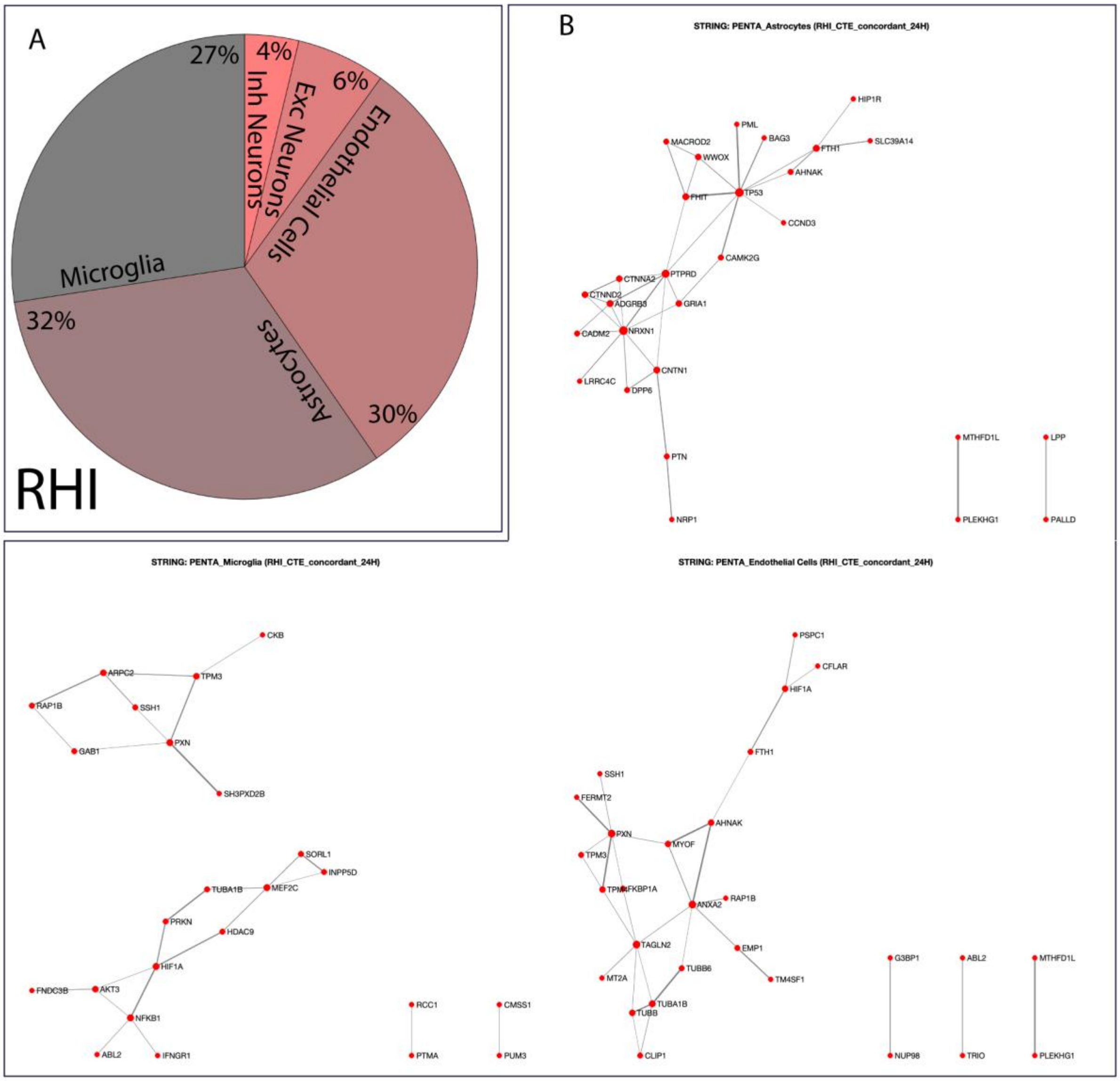


**Supplementary Figure 9.** Cell-type-resolved transcriptomic analyses were performed to characterize injury-associated pathway regulation and to identify dominant cellular contributors across human repetitive head injury (RHI) and chronic traumatic encephalopathy (CTE) datasets. **A** Pie chart showing the proportional contribution of major brain cell types to the RHI-associated gene set. **B** Network representations of significantly enriched pathways derived from CTE-associated genes, illustrating functional clustering and interconnectivity among injury-responsive biological processes. Nodes represent pathways and edges indicate shared gene membership.

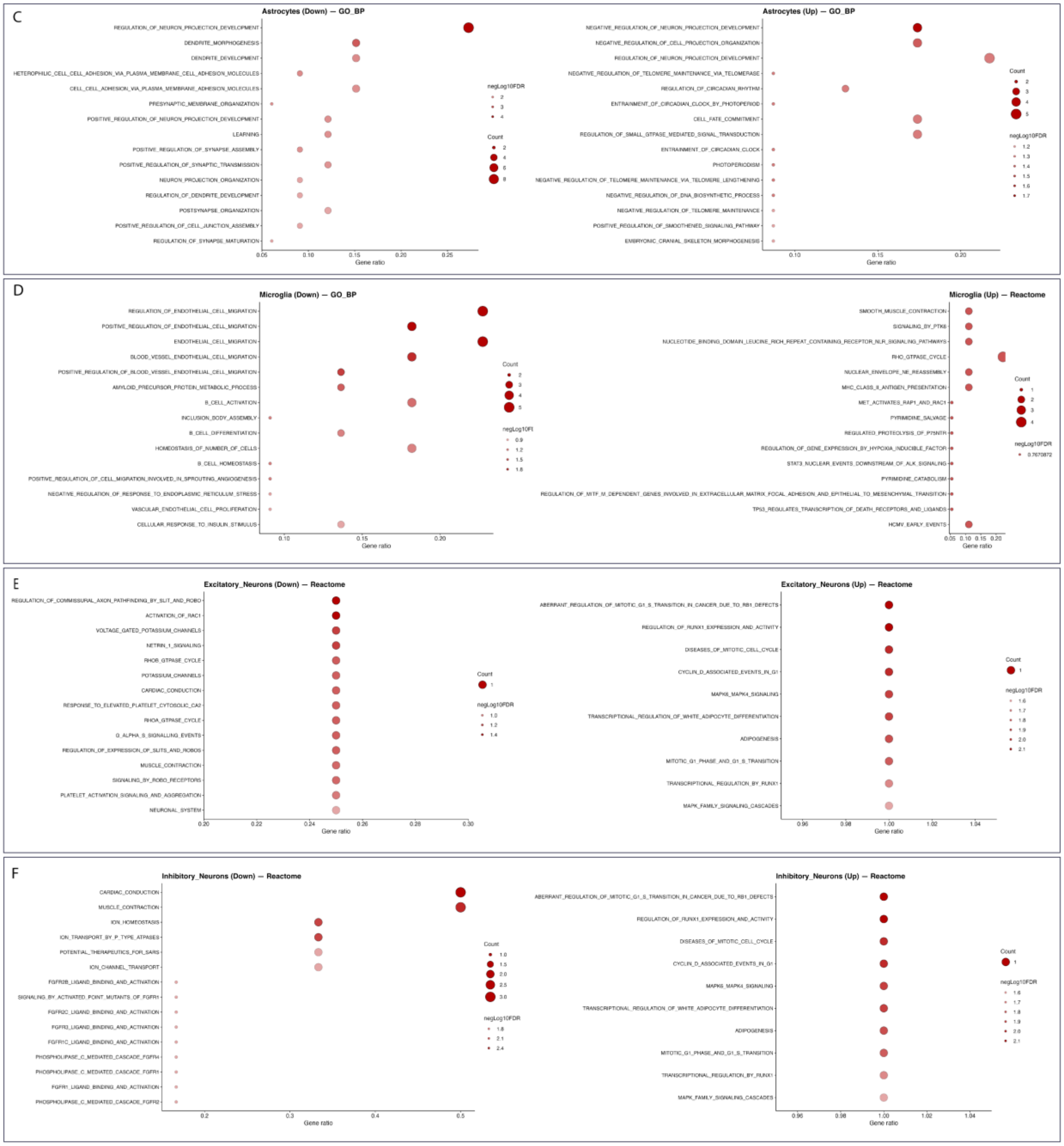


**Supplementary Figure 9. C** Dot plot visualization of Gene Ontology (GO) enrichment results for astrocytes in human CTE, shown separately for downregulated and upregulated pathways. Dot size represents the number of genes contributing to each pathway, and color intensity indicates statistical significance (−log10 adjusted *P* value). **D** Dot plot visualization of GO enrichment results for microglia in human CTE, separated into downregulated and upregulated pathways, highlighting enrichment of immune activation, inflammatory signaling, and cytoskeletal remodeling programs. **E** Reactome pathway enrichment analysis for excitatory neurons in human CTE,

illustrating suppression of synaptic organization, ion transport, and neuronal signaling pathways alongside activation of stress-associated programs.
**F** Reactome pathway enrichment analysis for inhibitory neurons in human CTE, revealing dysregulation of neurotransmission, metabolic, and cellular maintenance pathways linked to network imbalance.

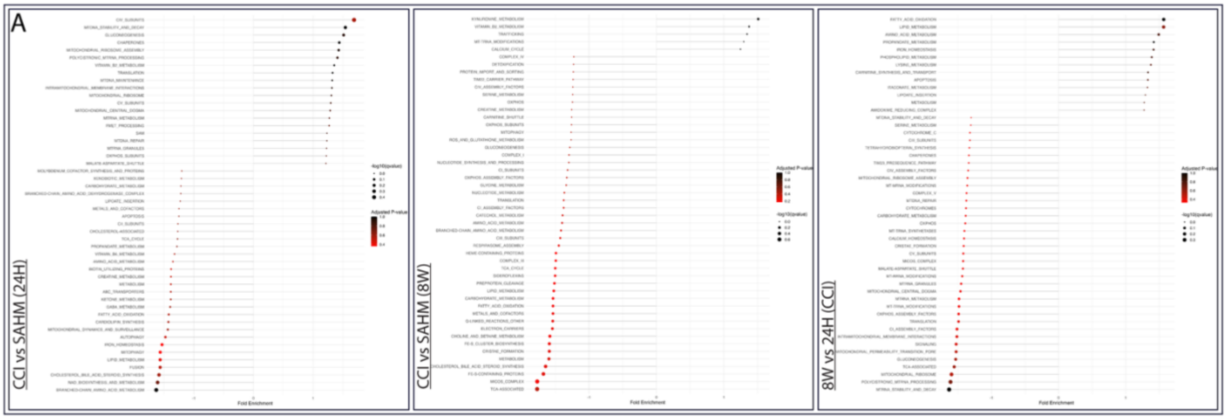


**Supplementary Figure 10**. Three mild repetitive injuries (1 mm-1 m/s) were applied to the NAM-triculture model with a 14-day interval. **A** RNA-Sequencing analysis comparison between CCI and SHAM 24 hours and 8 weeks post-injury. Comparison between 24 hours and 8 weeks among CCI.

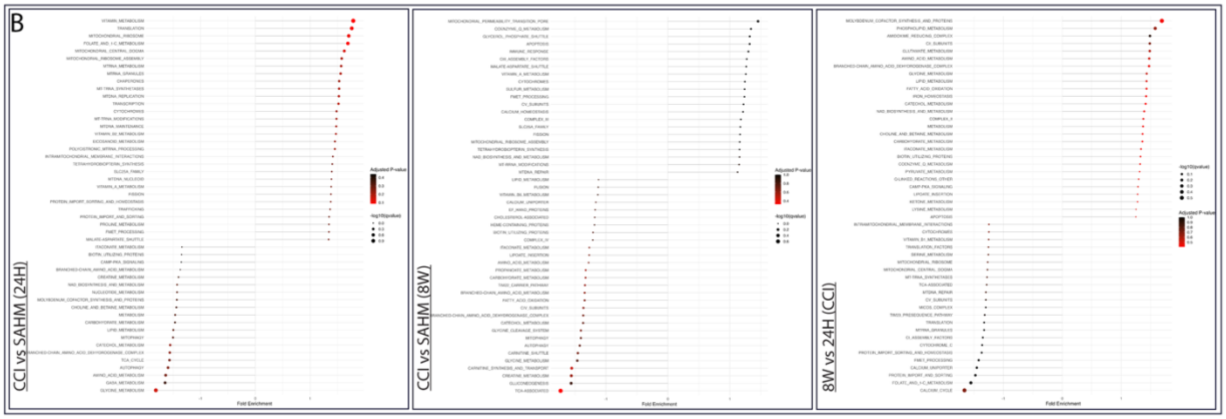


**Supplementary Figure 11.** Continuation of *Supplementary Figure 10***.** Three mild repetitive injuries (1 mm-1 m/s) were applied to the PENTA culture model with a 14-day interval. **B** RNA-Sequencing analysis comparison between CCI and SHAM 24 hours and 8 weeks post-injury. Comparison between 24 hours and 8 weeks among CCI.

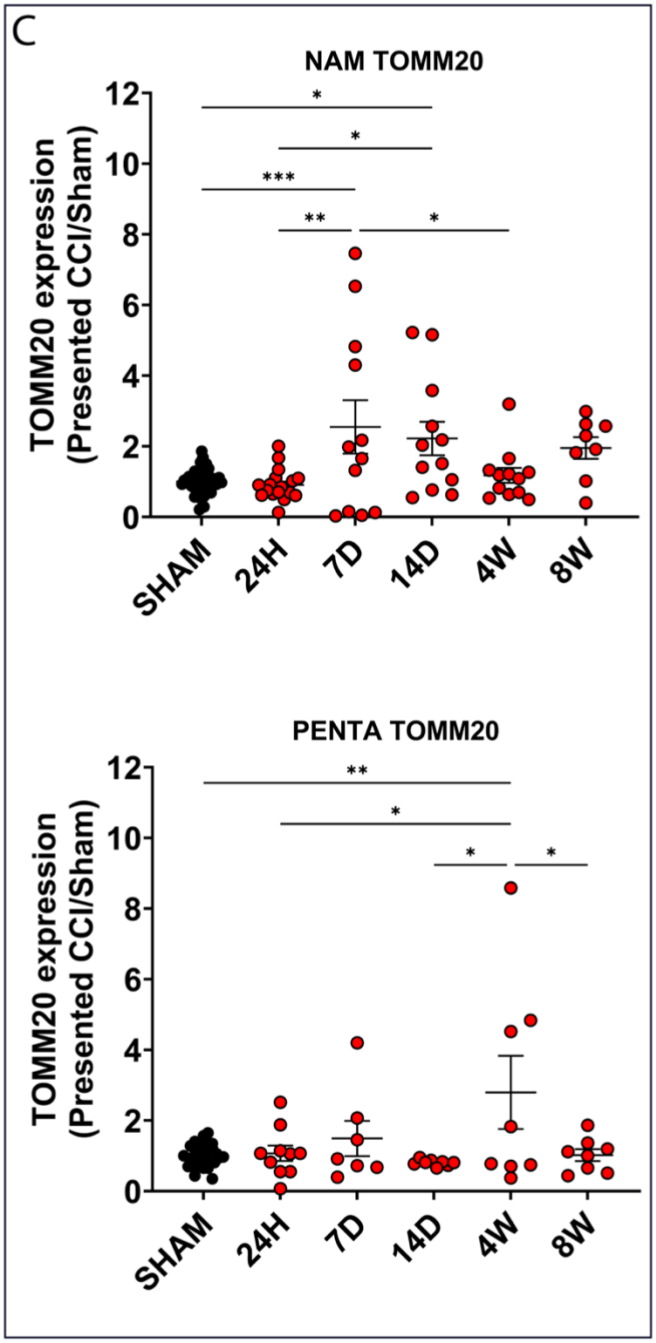


**Supplementary Figure 12.** Continuation of *Supplementary Figure 10***.** Intracellular markers – Western Blot quantification of TOMM 20. Data presented mean ± SEM of n = 6 – 12 scaffolds per condition. ***,** **, indicates a significant difference with p<0.05, p<0.01, respectively. A one-way ANOVA (analysis of variance) test was used to determine the difference between the control and experimental groups. The ROUT outlier analysis method was used to exclude statistical outliers. Experiments were replicated at least three times, unless specified otherwise. Intracellular markers for the 1W, 4W, and 8W time points were replicated twice.
